# Dynamic Interactions of Fully Glycosylated SARS-CoV-2 Spike Protein with Various Antibodies

**DOI:** 10.1101/2021.05.10.443519

**Authors:** Yiwei Cao, Yeol Kyo Choi, Martin Frank, Hyeonuk Woo, Sang-Jun Park, Min Sun Yeom, Chaok Seok, Wonpil Im

## Abstract

The spread of severe acute respiratory syndrome coronavirus 2 (SARS-CoV-2) presents a public health crisis, and the vaccines that can induce highly potent neutralizing antibodies are essential for ending the pandemic. The spike (S) protein on the viral envelope mediates human angiotensin-converting enzyme 2 (ACE2) binding and thus is the target of a variety of neutralizing antibodies. In this work, we built various S trimer-antibody complex structures on the basis of the fully glycosylated S protein models described in our previous work, and performed all-atom molecular dynamics simulations to get insight into the structural dynamics and interactions between S protein and antibodies. Investigation of the residues critical for S-antibody binding allows us to predict the potential influence of mutations in SARS-CoV-2 variants. Comparison of the glycan conformations between S-only and S-antibody systems reveals the roles of glycans in S-antibody binding. In addition, we explored the antibody binding modes, and the influences of antibody on the motion of S protein receptor binding domains. Overall, our analyses provide a better understanding of S-antibody interactions, and the simulation-based S-antibody interaction maps could be used to predict the influences of S mutation on S-antibody interactions, which will be useful for the development of vaccine and antibody-based therapy.

## INTRODUCTION

Since the outbreak of Coronavirus disease 2019 (COVID-19), the spread of severe acute respiratory syndrome coronavirus 2 (SARS-CoV-2) has infected over 150 million people and lead to 3.1 million deaths as of April, 2021. Due to the extremely high contagiousness of SARS-CoV-2 and lack of effective antiviral medicines, the neutralizing antibodies (NAbs) acquired through active or passive immunization have critical roles in preventing healthy individuals from infection and accelerating recovery of infected persons. Several SARS-CoV-2 vaccines have been authorized for emergency use, and the preliminary data show that they provide high protection rate^1-3^.

SARS-CoV-2 is an enveloped virus with a positive-sense single-stranded RNA genome^4^. The spike (S) trimer anchored in the viral envelope is a glycoprotein mediating the binding to human angiotensin-converting enzyme 2 (ACE2)^5-7^. S protein consists of multiple functional domains, including the receptor binding domain (RBD) that is responsible for interacting with ACE2. The RBDs on the top of S trimer are conformationally variable. In the so-called ‘closed state’, the RBDs lay flat with their receptor binding motifs (RBMs) occluded by the RBDs of neighboring protomers. The ‘open states’ are characterized by one or more uplifted RBDs, resulting in exposure of their RBMs (**Figure 1A**). Many NAbs have been isolated from the sera of recovered COVID-19 patients, and most of them target the RBD. The structures of S trimer in complex with various antibodies have been solved by cryogenic electron microscopy (cryo-EM), which provides valuable information of S-antibody interactions at near-atomic resolution^8-12^. However, such static structures may not include all the information that is necessary for understanding the mechanisms underlying antibody binding. In addition, many cryo-EM structures have missing residues in S RBDs and/or antibodies, and the glycans that can have significant influence on antibody binding are mostly not resolved. Therefore, molecular dynamics (MD) simulations based on well-refined initial structures with all missing portions properly modeled can provide a more comprehensive understanding of S-antibody interactions. As an RNA virus, SARS-CoV-2 has a relatively high mutation rate, which is a big challenge to the efficacy of antibodies and vaccines. In the past one and a half years, multiple variants of SARS-CoV-2 have appeared and are now circulating globally. For example, the B.1.1.7 variant that was initially detected in the UK has become a dominant strain in many countries^13^. By investigating the importance of each RBD residue in S-antibody interactions, we can evaluate the influence of the virus mutations, which can help to predict the efficacy of antibody against variant strains.

**Figure 1.**
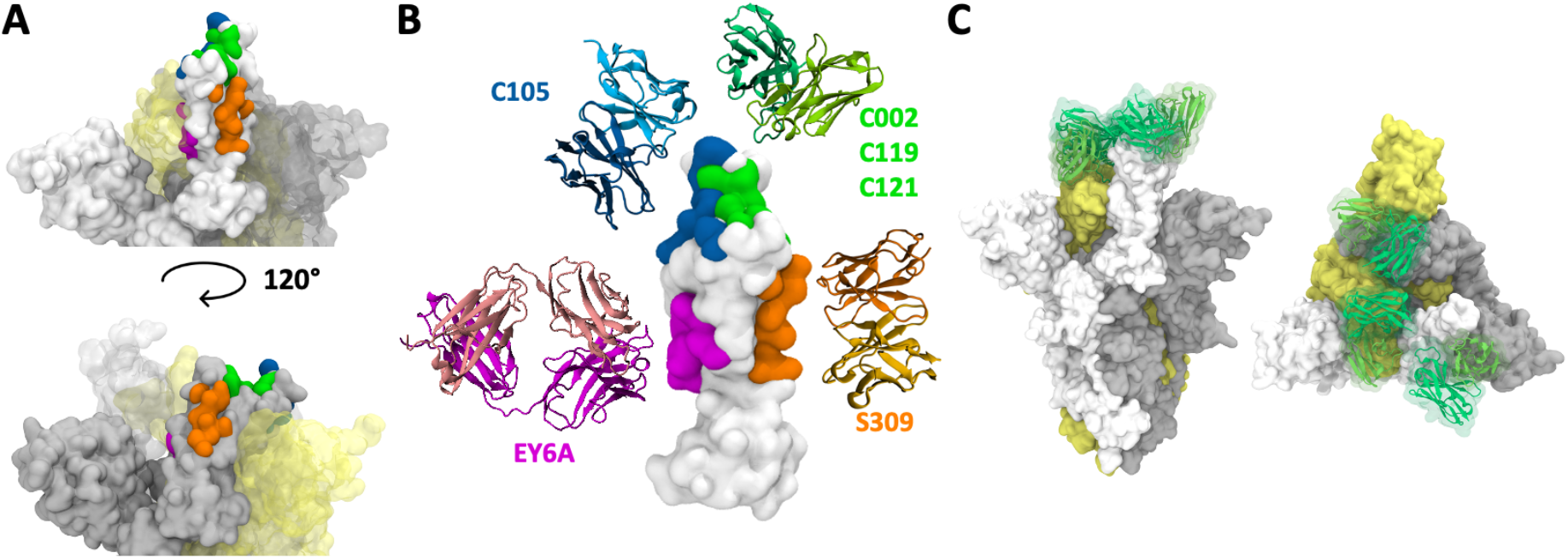
Overview of S RBD and various antibodies targeting different epitopes. (A) Open- and closed-state RBDs with epitopes marked by different colors. Three protomers of S protein are shown in white, gray, and yellow. (B) RBD epitopes and the corresponding antibodies studied in this work. The antibodies are arranged according to their binding sites. Antibodies C002, C119, and C121 share similar epitopes, and only one of them is shown in the figure. However, their epitope residues and binding poses are not completely identical. (C) A representative simulation system of S trimer with one open and two closed RBDs bound with three C119 antibodies (without water molecules and ions). All illustrations were created using Visual Molecular Dynamics (VMD)^14^.

In this work, we present all-atom MD simulations of a fully glycosylated S protein trimer in complex with various antibodies. We have selected antibodies that target different epitopes on the RBD (**Figure 1B**), and modeled the structures of S trimer bound with these antibodies (**Figure 1C**). 13 systems consisting of different antibodies and S trimers with different RBD open/closed states have been built and simulated (**Table 1**). For convenience, a one-letter symbol “O” or “C” is used to represent an open- or closed-state RBD, and thus antibody C002 bound to S trimer with open-closed-closed RBDs is denoted as “C002_OCC”. Except for antibody C105 whose epitope is not exposed when RBD is closed, we modelled all three RBDs bound with antibodies if the specific epitopes are not occluded and the modelled antibodies do not result in significant steric hinderance. (**Figure S1-S4)** Note that the reference PDB structures contain some free RBDs without bound antibodies. Our results provide insight into the dynamics of antibodies, contributions of residues for binding interactions, and roles of glycans in antibody binding, which provides a better understanding of S-antibody interactions.

**Table 1.**
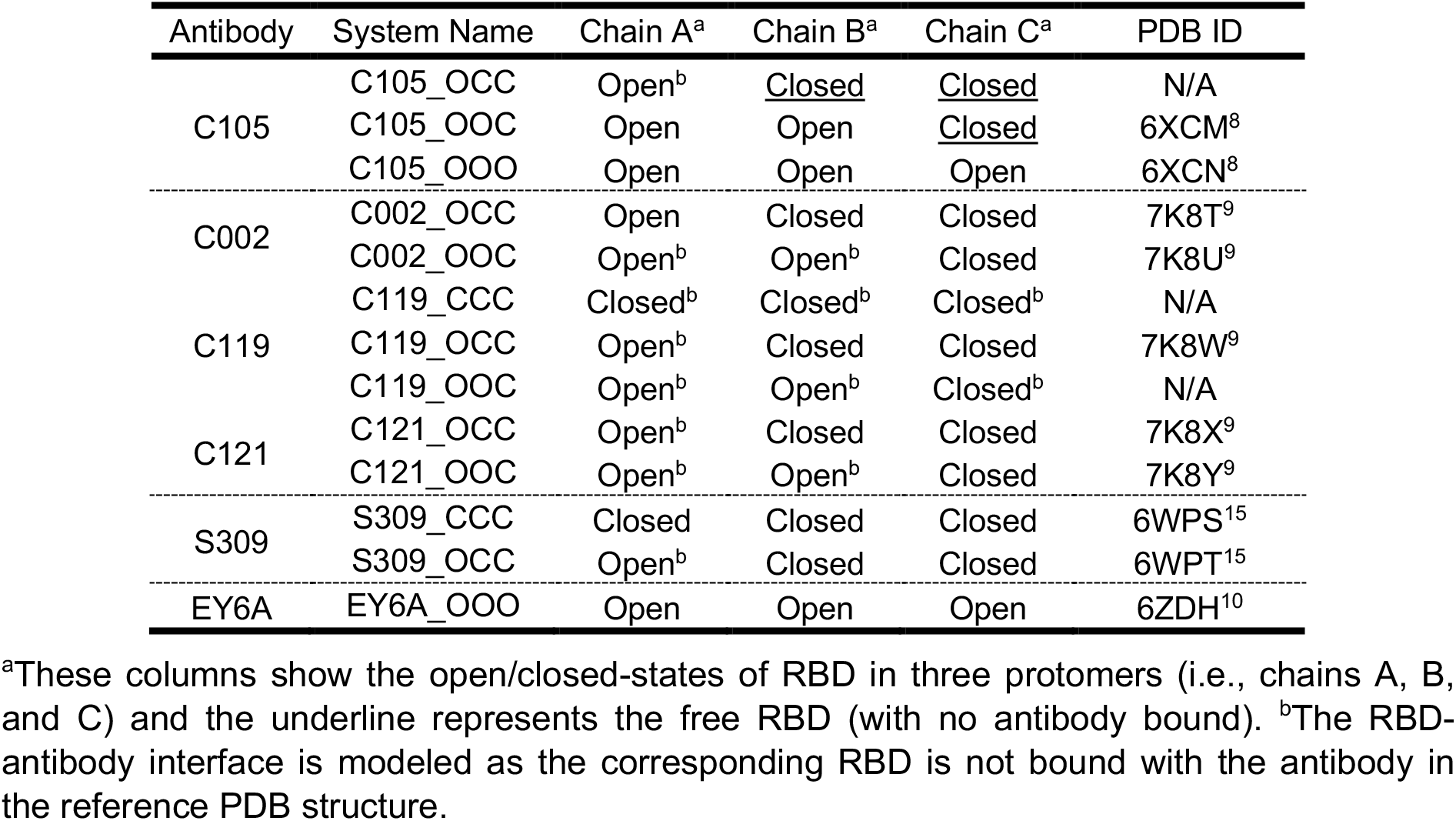
Simulation system information.

## METHODS

### Modeling of fully glycosylated SARS-CoV-2 S protein–antibody complex structures

As reported in our previous work^16-17^, we have modeled a fully glycosylated full-length SARS-CoV-2 S protein structure by using a combination of computational modeling tools including GALAXY protein modeling suite^18-20^ for building missing residues and domains, ISOLDE^21^ for model refinement against experimental density maps, and CHARMM-GUI *Glycan Reader & Modeler*^22-24^ and *Membrane Builder*^*25-28*^ for building glycans and a viral membrane. In this study, we truncated the heptad repeat linker, transmembrane domain, and cytoplasmic domain built by *ab initio* structure prediction and only kept the S1 subunit and part of S2 subunit. Before generating the S-antibody complex structures, we first removed all glycans to avoid bad contacts with antibodies. We extracted the portion of RBD-antibody from each cryo-EM structure of S trimer-antibody complex (**Table 1**), and superimposed it onto our S trimer structure by maximizing the overlap in RBD. For example, when building C121_OCC system, we used the cryo-EM structure (PDB id: 7K8X) as a reference. PDB 7K8X has two closed-state RBDs both bound with antibody C121, and one open-state unbound RBD. We simply extracted the portions of RBD-antibody from two closed chains in 7K8X and aligned them onto the closed chains in our S trimer structure. Since the open chain in 7K8X is unbound, we used the portion of RBD-antibody extracted from the closed chain to model the binding interface in the open chain with minor clashes. GALAXY protein modeling suite was then used to relax the structure to relieve the clashes. After the S trimer-antibody complex structure was generated, CHARMM-GUI *Glycan Reader & Modeler* was used to build 19 N-linked and 1 O-linked glycans onto each protomer using the same glycoforms as those in our previous model.

### Simulation details

In this study, the CHARMM36(m) force field was used for proteins^29^ and carbohydrates^30-32^. The TIP3P water model^33^ was utilized along with a 0.15 M KCl solution. The total number of atoms is approximately 1,250,000 (∼400,000 water molecules, ∼1,100 K^+^, and ∼1,100 Cl^-^), but the exact numbers differ between various systems. The van der Waals interactions were smoothly switched off over 10–12 Å by a force-based switching function^34^ and the long-range electrostatic interactions were calculated using the particle-mesh Ewald method^35^ with a mesh size of ∼1 Å. To avoid the overestimation of protein-protein interactions, a force field adjustment was made to enhance protein-water interactions^36-37^.

All simulations were performed using the input files generated by CHARMM-GUI^38-40^, and we used GROMACS 2018.6^41^ for both equilibration and production with the LINCS algorithm^42^. The temperature was maintained using a Nosé-Hoover temperature coupling method^43-44^ with a *τ*_t_ of 1 ps. For pressure coupling (1 bar), a semi-isotropic Parrinello–Rahman method^45-46^with a *τ*_p_ of 5 ps and a compressibility of 4.5×10^−5^ bar^−1^ was used. During the equilibration run, NVT (constant particle number, volume, and temperature) dynamics was first applied with a 1 fs time step for 250 ps. Subsequently, the NPT (constant particle number, pressure, and temperature) ensemble was applied with a 1-fs time step (for 125 ps) and with a 2-fs time step (for 1.5 ns). During the equilibration, positional and dihedral restraint potentials were applied, and their force constants were gradually reduced. The production run was performed with a 4 fs time step using the hydrogen mass repartitioning technique^47^ without any restraint potential. Each system shown in **Table 1** ran for 500 ns production time with about 20 ns/day using 768 CPU cores on NURION in the Korea Institute of Science and Technology Information. For comparison, we also ran 500 ns production of 12 S-only systems (3 for each of CCC, OCC, OOC, and OOO) with the same simulation protocols. All 4 S-only and 13 S-antibody simulation systems and trajectories are available in CHARMM-GUI COVID-19 archive (https://www.charmm-gui.org/docs/archive/covid19).

## RESULTS AND DISCUSSION

### Critical residues for antibody binding and potential influence of mutant variants

To identify critical residues for S-antibody interactions, we searched for all residue pairs consisting of one residue from S and the other from antibody, which make favorable interactions including hydrophobic interaction, *π*-*π* stacking interaction, hydrogen bond, and salt bridge. We processed 500 snapshots (every 1 ns) from the 500-ns simulation trajectory of each system, and calculated the frequency of interacting residue pairs.

**Figure 2** shows two representative cases of the antibodies bound to the RBDs. The first one is the interface of C105 bound to chain A in C105_OOC. There are 30 residues of RBD_A (RBD of chain A) that interact with the antibody in at least 10% of snapshots, and more than 10 of them keep their interactions with the antibody in at least 75% of snapshots (**Figure 2A**). Two of these critical residues are mutated in COVID-19 variants, which are K417N in South Africa variant (lineage B.1.351), K417T in Brazil variant (lineage P.1), and N501Y in UK/South Africa/Brazil variants. K417 has high frequency of interactions with Y33, Y52, and E96 from C105 V_H_ (variable domain of antibody heavy chain). As shown in **Figure 2C**, the positively-charged side chain of K417 can make a salt bridge with E96 or cation-*π* interaction with Y33 or Y52 depending on the conformation of these four residues, and such interactions are lost with K417N or K417T mutation. This implies that K417N/K417T will have significant effects on the interaction with C105, which is likely to reduce the efficacy of C105. N501 has high frequency of interactions with several residues from C105 V_L_ (variable domain of antibody light chain). Different from K417, N501 mostly interacts with C105 by the backbone atoms, and the side chain points toward the RBD itself. If a mutation does not significantly change the structure and dynamic of this local backbone, it should not be harmful to the binding of C105. However, N501Y introduces a new side chain with a larger size and the local backbone is a flexible loop. It is possible that the backbone conformational change is required to accommodate the side chain of Tyr. Therefore, it is possible that N501Y can be harmful to antibody binding.

**Figure 2.**
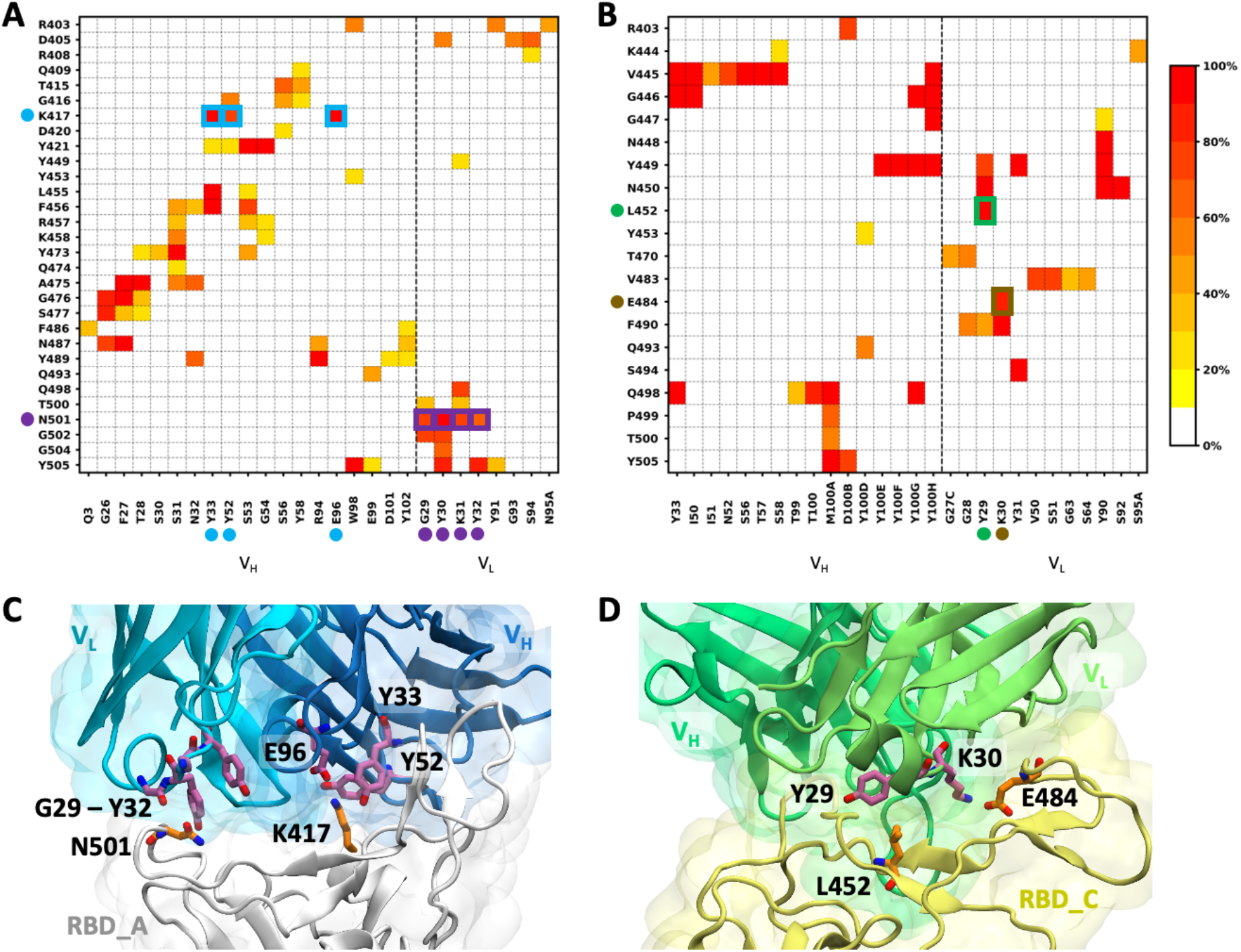
Frequency of interacting residue pairs between S RBD and antibody. Two representative examples are shown: (A) C105 bound to RBD_A in C105_OOC, and (B) C119 bound to RBD_C in C119_OCC. The x-axis labels the interacting residues of antibody, and the y-axis labels those of S protein. The color bar represents the frequency of interacting pairs observed in the simulation trajectory. We mark the point mutations occurring in COVID-19 variants and their interacting residues from antibodies by the dots with different colors if the residue pairs make favorable interactions in more than 60% of snapshots. Two snapshots with these residues highlighted are shown for (C) C105_OOC and (D) C119_OCC.

The second representative case is the interface of C119 bound to chain C in C119_OCC. There are more than 10 residues of RBD_C that continuously interact with C119 V_H_ and/or V_L_ in the simulation trajectory (**Figure 2B**). Two critical residues involved in COVID-19 variants, L452R and E484K, occur simultaneously as a double mutation in India variant (lineage B.1.617). E484K also appears in South Africa and Brazil variants as well as some sequences but not all in UK variant, and L452R appears in California variant (lineages B.1.427/B.1.429). As shown in **Figure 2D**, the hydrophobic side chain of L452 continuously interacts with the aromatic side chain of Y29 from C119 V_L_. With the L452R mutation, the positively charged side chain can still interact with the aromatic ring (i.e., cation-*π* interaction), hence it may not disturb the binding of C119. In contrast, E484 has a salt bridge with K30 from C119 V_L_. The E484K mutation not only breaks the salt bridge but also forms an unfavorable contact of two positively charged groups. Therefore, E484K may seriously impair the efficacy of antibody C119.

The interacting residue pairs of the all systems are shown in **Figure S5-S10**. In summary, antibody C105 is likely to be sensitive to mutation K417N/K417T as salt bridge and cation-*π* interaction involving K417 is frequently observed in all cases except chain A of C105_OOO. However, the influence of N501Y is uncertain and the interaction between N501 and antibody is only observed in chain A of C105_OOC. Antibodies C002, C119, and C121 have frequent interactions with L452 and/or E484, which is observed in all cases except chain A of C121_OCC. Since all variants of current concern contain at least one of two mutations, it remains a question whether these three antibodies are effective enough to the variants. Antibodies S309 and EY6A target different epitopes that do not contain the mutant residues, and hopefully, they would not be affected by the mutations in COVID-19 variants. The maps of interacting pairs derived from our simulation results highlight the critical residues from RBD and their interacting residues from antibodies. By considering the physicochemical properties of wild-type and mutant amino acids as well as their possible interactions with the antibody residues, we are able to predict the potential impacts of S mutations on antibody binding.

### Antibodies can have two binding interfaces with RBDs from different protomers

The antibodies C002, C119, and C121 bind to similar but not identical epitopes, located around the green region shown in **Figure 1**, with different interaction patterns. When these antibodies bind to a closed-state RBD (e.g., RBD_C), they can also have a secondary binding interface with the next RBD in the anti-clockwise direction from top view (e.g., RBD_A). To examine the existence of such a secondary binding interface for other systems, we performed the analysis of interacting residue pairs with consideration of both RBD containing the primary interface and the neighboring RBD that may contain the secondary interface (**Figure S5-S10**). Except for C002, C119, and C121, all other antibodies have only one primary binding interface. C002, C119, and C121 have a secondary binding interface regardless of whether the neighboring RBD is open or closed, but the secondary interface has more extensive interacting residue pairs when the neighboring RBD is open (**Figure S6-S8**).

**Figure 3** shows two representative cases of the antibodies interacting with two RBDs. The first is C121 bound to closed-state RBD_C in C121_OCC. C121 has a secondary binding interface with the neighboring open-state RBD_A. **Figure 3A** shows that V_H_ makes the major contribution to RBD_C binding. In addition, there are multiple residues from both C121 V_H_ and V_L_ that continuously interact with the residues from RBD_A. This suggests that C121 binds to S protein through multivalent interactions which can lead to enhancement of binding affinity. As shown in **Figure 3C**, both complementarity-determining region 3 (CDR3) of V_H_ (red circle) and CDR1/CRD2 of V_L_ (purple circle) are involved in the secondary binding interface. When C121 binds to RBD_B (primary) and RBD_C (secondary) that are both closed, the secondary interface consists of only a couple of residues from CDR3 of V_H_, but two of them (V106 and L107) continuously interact with RBD_C (**Figure S8A** middle).

**Figure 3.**
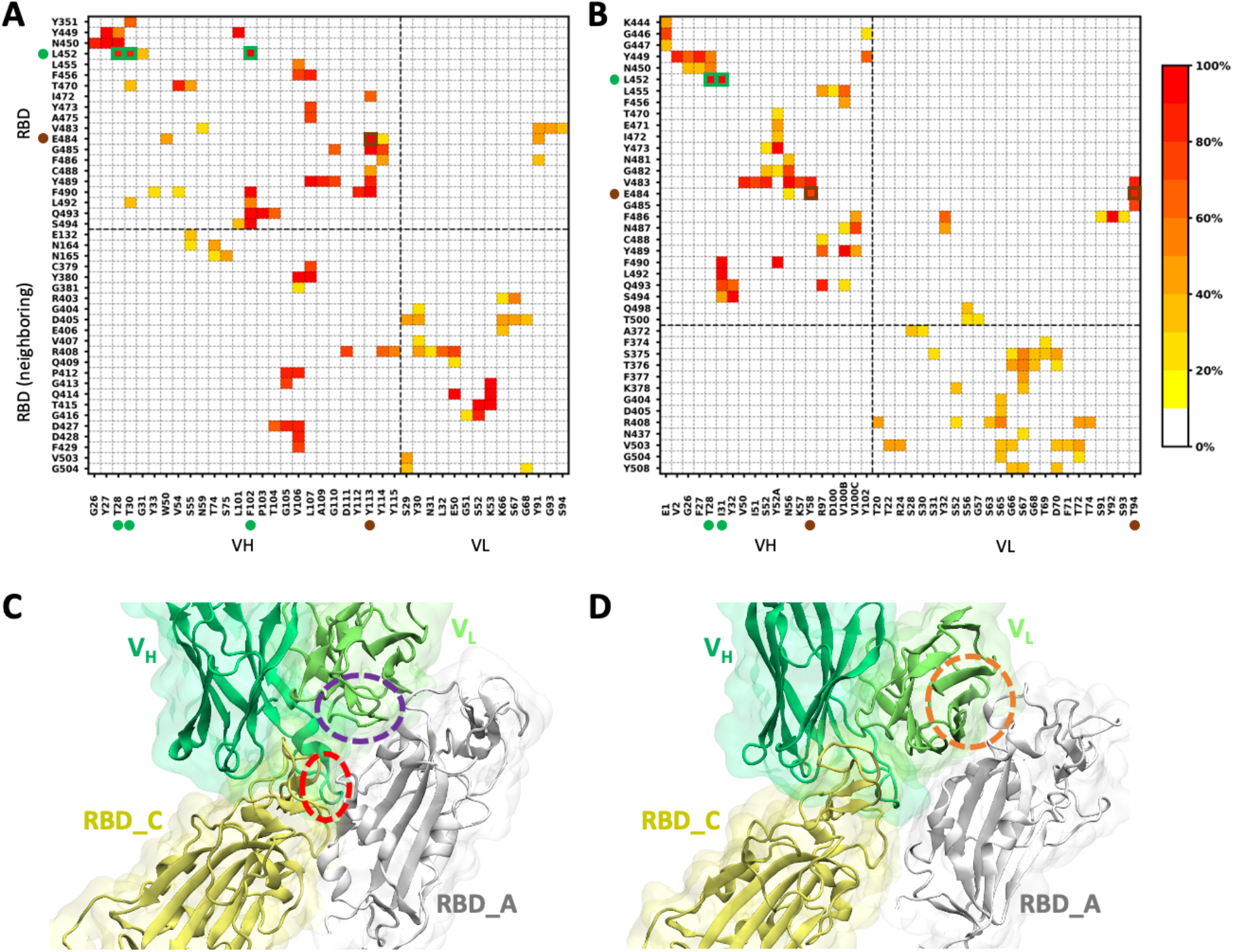
Frequency of interacting residue pairs in both primary and secondary binding interfaces. Two representative examples are shown: (A) C121 bound to RBD_C (primary) and RBD_A (secondary) in C121_OCC, and (B) C002 bound to RBD_C (primary) and RBD_A (secondary) in C002_OCC. The x-axis labels the interacting residues of antibody, and the y-axis labels those of S protein. The color bar represents the frequency of interacting pairs observed in the simulation trajectory. We also mark the point mutations occurring in COVID-19 variants and their interacting residues from antibodies by the dots with different colors if the residue pairs make favorable interactions in more than 60% of snapshots. Two snapshots illustrating the binding poses of two interfaces are shown for (C) C121_OCC and (D) C002_OCC. The antibody residues involved in the secondary interfaces are encircled.

The second representative case is C002 bound to closed-state RBD_C (primary) and open-state RBD_A (secondary) in C002_OCC. Although multiple residues from V_L_ are involved in the secondary interface, none of the interacting residue pairs has a frequency over 60%, indicating that the residue contacts in the secondary interface keep varying with the movements of V_L_ and RBD_A (**Figure 3B**). As shown in **Figure 3D**, the antibody residues involved in the secondary interface (orange circle) are not within any CDR of V_L_. Therefore, the binding interaction between C002 and RBD_C/RBD_A is not multivalent, and the secondary interface appears to exist simply because the binding pose of C002 in the primary interface and the relative position of two RBDs leads to such fortuitous contacts.

### RBD-antibody interfaces are stable in all systems

Although there exist cryo-EM structures for each antibody, many of them have only one or two antibodies bound to RBD(s). In this study, we aimed to build the S-antibody complex systems with all three RBDs bound with antibodies, except for C105 whose epitope is not exposed when RBD is closed. Therefore, we needed to check whether binding interfaces are stable considering the mutual influence between multiple antibodies. For each system, we measured the minimum distance from V_H_/V_L_ to RBD, and number of interacting residue pairs between V_H_/V_L_ and RBD (**Figure S11-S16**). For the antibodies with primary and secondary interfaces, we considered only the primary interfaces. All antibodies stably stay in the binding sites and continuously interact with the epitopes on RBDs (at least during the current simulation time). We observed three different binding modes; (i) both V_H_ and V_L_ make similar contributions to RBD binding (**Figure 4A**), (ii) V_H_ makes the major contribution to RBD binding, but V_L_ continuously has contacts with RBD (**Figure 4B**), and (iii) V_H_ makes the major contribution to RBD binding and V_L_ can move away from the RBD (**Figure 4C**). A summary of binding modes in each system is shown in **Table 2**. Clearly, S-antibody binding modes are versatile and specific to individual antibodies and RBD’s open/closed states.

**Table 2.**
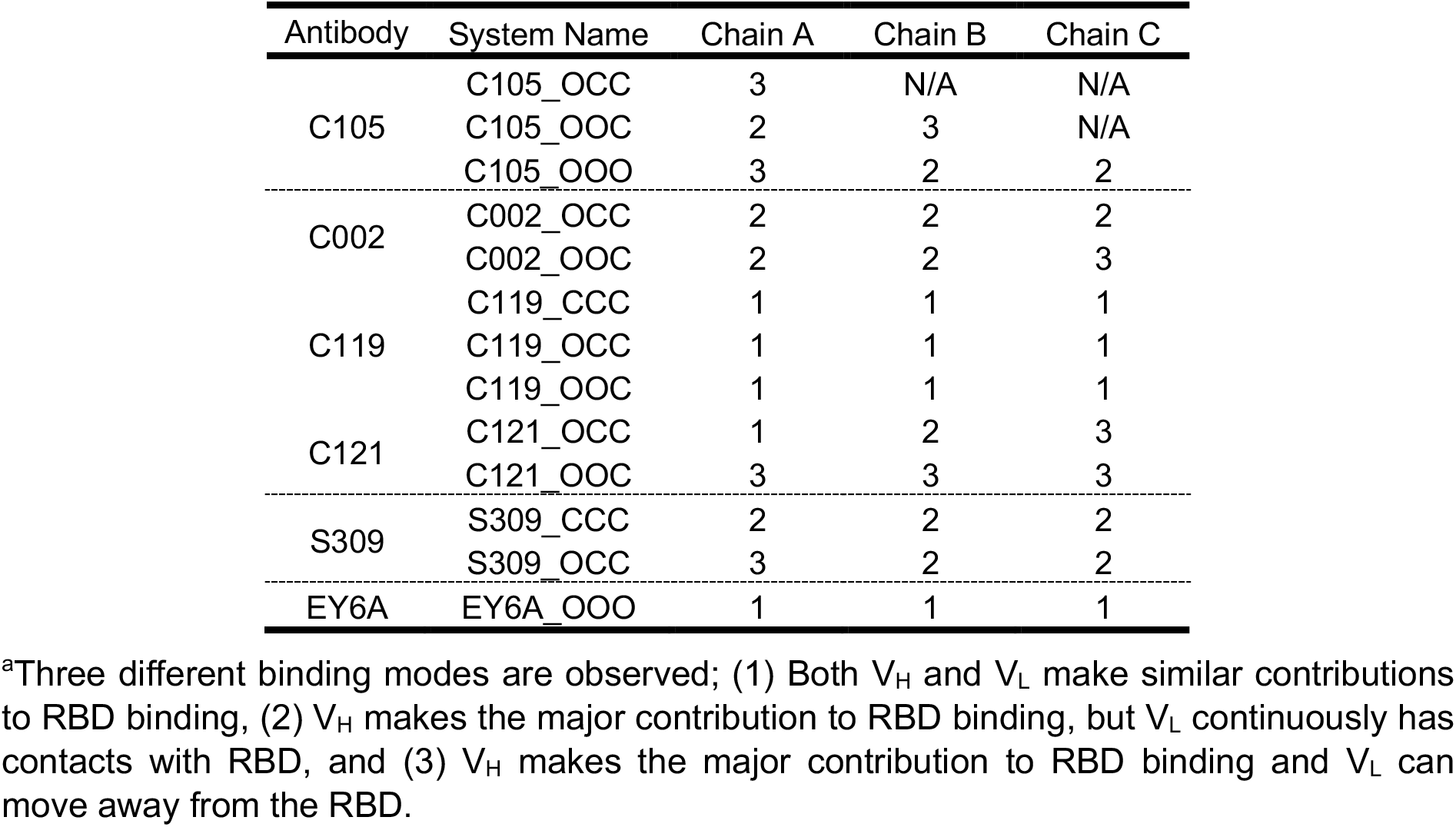
Antibody binding mode^a^.

**Figure 4.**
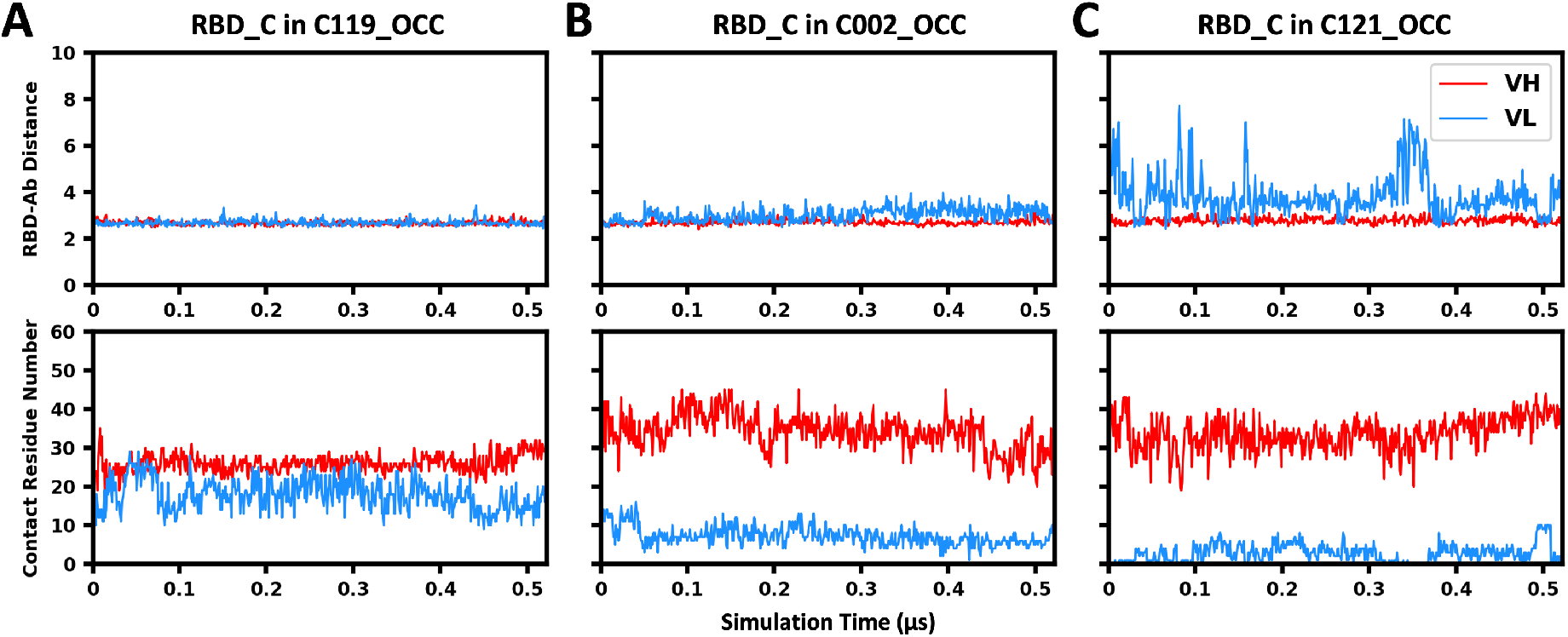
RBD-antibody minimum distance and contact residue number in representative binding interfaces. (A) C119 bound to RBD_C in C119_OCC, (B) C002 bound to RBD_C in C002_OCC, and (C) C121 bound to RBD_C in C121_OCC.

### Glycan flexibility and its roles in antibody binding

S protein is highly glycosylated and it has been shown that the glycans can have important roles in protein stability, RBD open-closed state transition, and interactions with ACE2 and antibodies^48-50^. In our previous work^17^, we aligned the PDB structures of RBD-antibody complexes onto the simulation trajectories of fully-glycosylated S trimers, and investigated the influences of glycans on antibody binding by measuring the extent of clashes between glycans and superimposed antibodies. In this work, we explored the same question by comparing the glycan flexibility in S-only and S-antibody complex systems. Four glycans attached to N122, N165, N331, and N343 are close to the epitopes marked with green and orange in **Figure 1**. All of them could possibly influence antibody binding depending on the glycan conformation and open/closed state of RBD. Among these four glycans, N343 glycan is attached to the residue within the orange epitope, and N165 glycan continuously interacts with the green epitope, hence they are more likely to be involved in antibody binding activity. We measured the φ/ψ angles of the first three glycosidic linkages from the reducing terminal (i.e., the linkages in Manβ1-4GlcNAcβ1-4 GlcNAcβ1-Asn) for N343 in S309_CCC and N165 glycan in C119_CCC, as well as these two glycans in the S-only system with three closed-state RBDs. The dihedral angles of GlcNAcβ1-Asn linkage are defined by φ: O_5_(1)-C_1_(1)-N_δ_(2)-C_γ_(2) and ψ: C_1_(1)-N_δ_(2)-C_γ_(2)-C_β_(2), where (1) is GlcNAc and (2) is Asn. The dihedral angles of Manβ1-4GlcNAc and 4GlcNAcβ1-4 GlcNAc linkages are defined by φ: O_5_(1)-C_1_(1)-O_4_(2)-C_4_(2) and ψ: C_1_(1)-O_4_(2)-C_4_(2)-C_3_(2), where (1) is the non-reducing terminal unit and (2) is the reducing terminal unit.

For illustration, we superimposed the N343 glycan in each snapshot onto the initial structures and the resulting structures of S-only and S-antibody systems are shown in **Figure 5A. Figure 5B** shows the distributions of dihedral angles in N343 glycosidic linkages, and the glycans attached to RBD_A, RBD_B, and RBD_C are shown in different colors. By comparing two rows, it becomes evident that the conformational space explored by N343 glycan in the S-only system is similar to that in the S-antibody system. Only two minor regions that are explored in the S-only systems (labeled by red arrows) are not explored in the S-antibody systems. In the S-only system, N343 glycan interacts with the closed-state RBD on the neighboring chain in most of the simulation time. This interaction keeps N343 glycan pointing towards the left direction (**Figure 5A**), which does not disturb the binding of antibody S309. On the contrary, the core portion of the glycan can interact with the antibody, serving as part of the epitope. Only very occasionally, it switches to the other direction and shields the binding interface for the antibody. Therefore, N343 glycan facilitates the binding of antibody S309 rather than blocking such interaction, which is consistent with the conclusion from our previous work.

**Figure 5.**
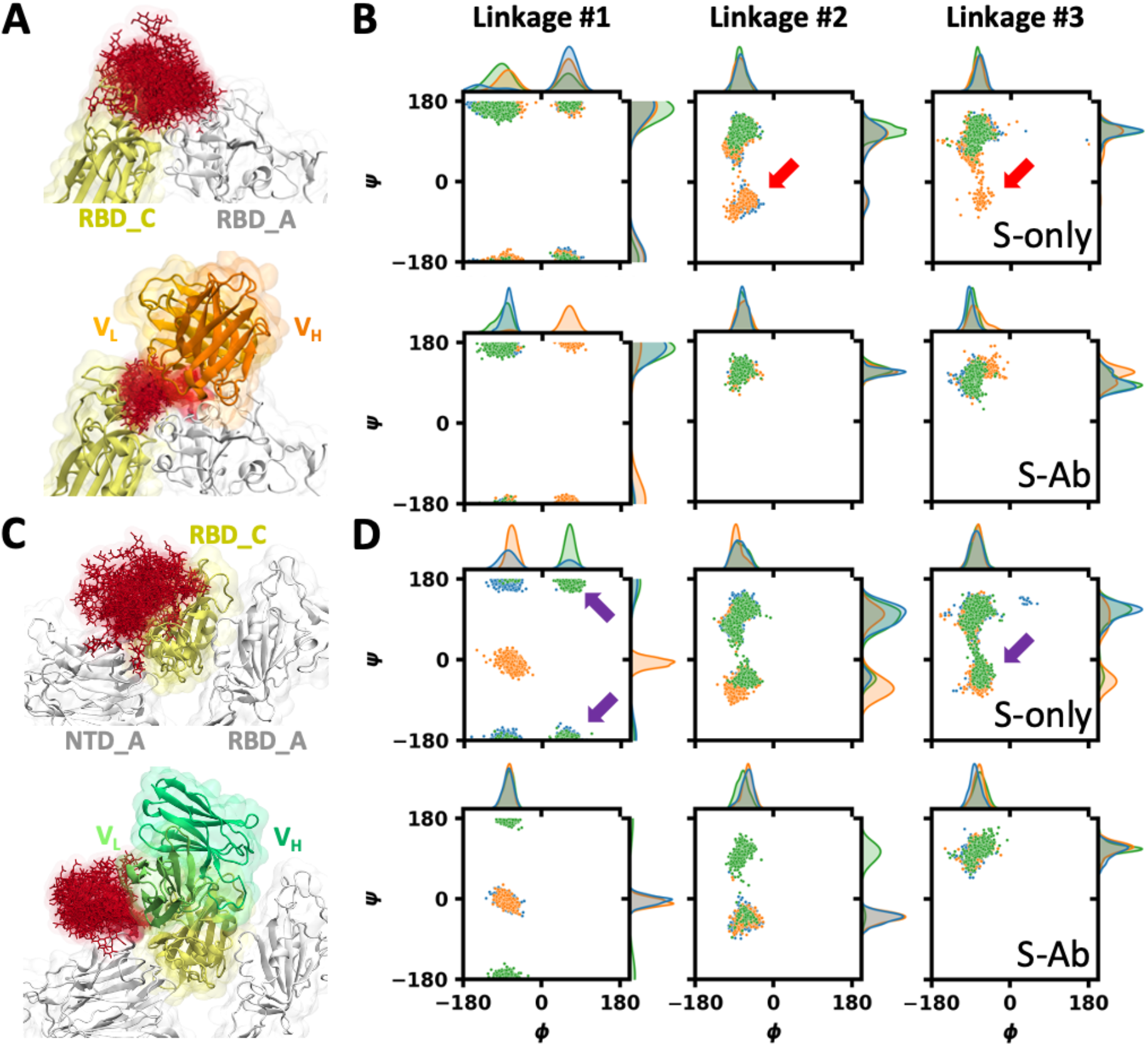
Glycan flexibility in S-only and S-antibody complex systems. The distributions of N-glycan structures are shown for (A) N343 glycan in S309_CCC and (C) N165 glycan in C119_CCC, where the glycan (red) in each snapshot is superimposed onto the initial structure by aligning the local backbone consisting of the glycosylation site and two neighboring residues. The corresponding distributions of φ/ψ angles in the first three glycosidic linkages from the reducing terminus are plotted for (B) N343 glycan and (D) N165 glycan. The glycans attached to three protomers (chains A, B, and C) are shown in blue, orange, and green, respectively.

**Figure 5D** shows the comparison of dihedral distributions of N165 glycan in S-only and S-antibody systems. There are two major regions (labeled by purple arrows) that are explored in the S-only system but not in S-antibody system. As shown in the superimposed glycan conformations (**Figure 5C**), when the antibody exists, the glycan is forced to point towards the left direction. Note that we built the S-antibody system by first generating S trimer-antibody complex and then modeling the glycans, and consequently we observed that the antibody restricted the glycan motion. However, if we compare this with the simulation of the unbound S glycoprotein, it becomes clear that N165 glycan occupied the binding interface for antibody C119 in about half of the simulation time and moved out of that region in the remaining time. These indicate that N165 glycan does not always block the antibody binding, but the existence of this glycan makes it more difficult for the antibody to get access to the epitope.

### Effects of antibody binding on RBD motion and flexibility

To investigate whether antibody binding has significant influences on RBD motion, we measured two structural features: the RBD-NTD (N-terminal domain) distance (*d*) defined by the minimum distance between RBD and NTD, and the RBD orientation angle (*θ*) defined by three points on RBD and S-trimer central axis (**Figure S17**). *θ* reflects the open/closed state of RBD, and *d* reflects the size of U-shape pocket that accommodates the neighboring closed-state RBD. It is expected that antibody binding could stabilize the S trimer and make the RBD open/closed state transition more difficult, particularly for the antibodies having two interfaces with different protomers. For all-closed and one-open S trimer, the RBDs are generally stable and show very limited motions even in the S-only systems, hence antibody binding has generally no influence on RBD orientation and motion (during the current simulation time). Additionally, it becomes evident that it would require a much longer simulation time to explore the RBD open-closed state transition in S-only and S-antibody systems. For two-open and all-open S trimer, the open-state RBDs move freely and exhibit huge motions even on the 500-ns simulation time scale, which is not affected by antibody binding either.

In addition to RBD motion, we also measured the root-mean-square fluctuation (RMSF) of RBD in S-only and S-antibody systems (**Figure S18-S21**). Though there exist differences between different protomers, the open-state RBDs generally have higher RMSF than the closed-state RBDs. The antibodies target both highly-flexible regions and stable regions. There are two large peaks around residues 445 and 480, which are two loops on the top of RBD. They become much more stable upon antibody binding, particularly when these regions are within the epitopes of antibodies (i.e., C002, C119, and C121).

## CONCLUSIONS

In this work, we have presented a modeling and simulation study of fully glycosylated S protein in complex with various antibodies. The frequency of interacting residue pairs between S and antibody reveals the interaction patterns and the critical S protein residues contributing to antibody binding, which enables us to predict possible effects of mutations in SARS-CoV-2 variants. Such analyses also made it possible to identify the antibodies that bind the S trimer through multivalent interactions. Comparison of glycan conformational flexibility in S-only and S-antibody complex systems reveals whether a specific glycan disturbs antibody binding. During the current limited simulation time, we could not observe that antibodies had any significant influence on RBD open-closed state transition, but the RMSF of RBD is reduced due to antibody binding. Together, this study provides richer insight into the S-antibody interactions that is not available from the static cryo-EM structures, and we hope that our findings are useful for the development of vaccines and antibody-based therapeutic agents.

## ACKNOWLEDGMENT

This study was supported in part by grants from NIH GM126140, GM138472, NSF OAC-1931343, DBI-2011234, DBI-1660380, and MCB-1810695, a Friedrich Wilhelm Bessel Research Award from the Humboldt Foundation (WI), National Research Foundation of Korea grants (2019M3E5D4066898 and 2016M3C4A7952630) funded by the Korea government (CS), the National Supercomputing Center with supercomputing resources including technical support (KSC-2020-CRE-0089 and KSC-2020-CRE-0094) (MSY and CS), and COVID-19 HPC Consortium project BIO200063 (https://covid19-hpc-consortium.org/).

## SUPPORTING INFORMATION

Figure S1. Illustration of C105_OCC, C105_OOC and C105_OOO. Figure S2. Illustration of C119_CCC, C119_OCC and C119_OOC. Figure S3. Illustration of S309_CCC, and S309_OCC. Figure S4. Illustration of EY6A_OOO. Figure S5. The frequency of interacting residue pairs in C105 systems. Figure S6. The frequency of interacting residue pairs in C002 systems. Figure S7. The frequency of interacting residue pairs in C119 systems. Figure S8. The frequency of interacting residue pairs in C121 systems. Figure S9. The frequency of interacting residue pairs in S309 systems. Figure S10. The frequency of interacting residue pairs in EY6A_OOO. Figure S11. RBD-antibody distance and contact residue number in C105 systems. Figure S12. RBD-antibody distance and contact residue number in C002 systems. Figure S13. RBD-antibody distance and contact residue number in C119 systems. Figure S14. RBD-antibody distance and contact residue number in C121 systems. Figure S15. RBD-antibody distance and contact residue number in S309 systems. Figure S16. RBD-antibody distance and contact residue number in EY6A_OOO. Figure S17. NTD-RBD distance and RBD orientation in S-only and S-antibody complex systems. Figure S18. RMSF of RBD in S-only and S-antibody complex systems with all three RBDs closed. Figure S19. RMSF of RBD in S-only and S-antibody complex systems with one RBD open and two RBDs closed. Figure S20. RMSF of RBD in S-only and S-antibody complex systems with two RBDs open and one RBD closed. Figure S21. RMSF of RBD in S-only and S-antibody complex systems with all three RBDs open. This material is available free of charge via the Internet at http://pubs.acs.org.

## NOTES

The authors declare no competing financial interest.

## REFERENCES

1. Polack, F. P., Thomas, S. J., Kitchin, N., Absalon, J., Gurtman, A., Lockhart, S., Perez, J. L., Pérez Marc, G., Moreira, E. D., Zerbini, C., Bailey, R., Swanson, K. A., Roychoudhury, S., Koury, K., Li, P., Kalina, W. V., Cooper, D., Frenck, R. W., Hammitt, L. L., Türeci, Ö., Nell, H., Schaefer, A., Ünal, S., Tresnan, D. B., Mather, S., Dormitzer, P. R., Şahin, U., Jansen, K. U., Gruber, W. C. N. Engl. J. Med. 2020, 383, 2603–2615.

2. Baden, L. R., El Sahly, H. M., Essink, B., Kotloff, K., Frey, S., Novak, R., Diemert, D., Spector, S. A., Rouphael, N., Creech, C. B., McGettigan, J., Khetan, S., Segall, N., Solis, J., Brosz, A., Fierro, C., Schwartz, H., Neuzil, K., Corey, L., Gilbert, P., Janes, H., Follmann, D., Marovich, M., Mascola, J., Polakowski, L., Ledgerwood, J., Graham, B. S., Bennett, H., Pajon, R., Knightly, C., Leav, B., Deng, W., Zhou, H., Han, S., Ivarsson, M., Miller, J., Zaks, T. N. Engl. J. Med. 2021, 384, 403–416.

3. Dagan, N., Barda, N., Kepten, E., Miron, O., Perchik, S., Katz, M. A., Hernán, M. A., Lipsitch, M., Reis, B., Balicer, R. D. N. Engl. J. Med. 2021, 384, 1412–1423.

4. Lai, M. M. C., Cavanagh, D. Adv. Virus Res. 1997, 48, 1–100.

5. Letko, M., Marzi, A., Munster, V. Nat. Microbiol. 2020, 5, 562–569.

6. Hoffmann, M., Kleine-Weber, H., Schroeder, S., Krüger, N., Herrler, T., Erichsen, S., Schiergens, T. S., Herrler, G., Wu, N.-H., Nitsche, A., Müller, M. A., Drosten, C., Pöhlmann, S. Cell 2020, 181, 271-280.e8.

7. Wrapp, D., Wang, N., Corbett, K. S., Goldsmith, J. A., Hsieh, C.-L., Abiona, O., Graham, B. S., McLellan, J. S. Science 2020, 367, 1260–1263.

8. Barnes, C. O., West, A. P., Huey-Tubman, K. E., Hoffmann, M. A. G., Sharaf, N. G., Hoffman, P. R., Koranda, N., Gristick, H. B., Gaebler, C., Muecksch, F., Lorenzi, J. C. C., Finkin, S., Hägglöf, T., Hurley, A., Millard, K. G., Weisblum, Y., Schmidt, F., Hatziioannou, T., Bieniasz, P. D., Caskey, M., Robbiani, D. F., Nussenzweig, M. C., Bjorkman, P. J. Cell 2020, 182, 828–842.e16.

9. Barnes, C. O., Jette, C. A., Abernathy, M. E., Dam, K.-M. A., Esswein, S. R., Gristick, H. B., Malyutin, A. G., Sharaf, N. G., Huey-Tubman, K. E., Lee, Y. E., Robbiani, D. F., Nussenzweig, M. C., West, A. P., Bjorkman, P. J. Nature 2020, 588, 682–687.

10. Zhou, D., Duyvesteyn, H. M. E., Chen, C.-P., Huang, C.-G., Chen, T.-H., Shih, S.-R., Lin, Y.-C., Cheng, C.-Y., Cheng, S.-H., Huang, Y.-C., Lin, T.-Y., Ma, C., Huo, J., Carrique, L., Malinauskas, T., Ruza, R. R., Shah, P. N. M., Tan, T. K., Rijal, P., Donat, R. F., Godwin, K., Buttigieg, K. R., Tree, J. A., Radecke, J., Paterson, N. G., Supasa, P., Mongkolsapaya, J., Screaton, G. R., Carroll, M. W., Gilbert-Jaramillo, J., Knight, M. L., James, W., Owens, R. J., Naismith, J. H., Townsend, A. R., Fry, E. E., Zhao, Y., Ren, J., Stuart, D. I., Huang, K.-Y. A. Nat. Struct. Mol. Biol. 2020, 27, 950–958.

11. Tortorici, M. A., Beltramello, M., Lempp, F. A., Pinto, D., Dang, H. V., Rosen, L. E., McCallum, M., Bowen, J., Minola, A., Jaconi, S., Zatta, F., De Marco, A., Guarino, B., Bianchi, S., Lauron, E. J., Tucker, H., Zhou, J., Peter, A., Havenar-Daughton, C., Wojcechowskyj, J. A., Case, J. B., Chen, R. E., Kaiser, H., Montiel-Ruiz, M., Meury, M., Czudnochowski, N., Spreafico, R., Dillen, J., Ng, C., Sprugasci, N., Culap, K., Benigni, F., Abdelnabi, R., Foo, S.-Y. C., Schmid, M. A., Cameroni, E., Riva, A., Gabrieli, A., Galli, M., Pizzuto, M. S., Neyts, J., Diamond, M. S., Virgin, H. W., Snell, G., Corti, D., Fink, K., Veesler, D. Science 2020, 370, 950–957.

12. Liu, L., Wang, P., Nair, M. S., Yu, J., Rapp, M., Wang, Q., Luo, Y., Chan, J. F. W., Sahi, V., Figueroa, A., Guo, X. V., Cerutti, G., Bimela, J., Gorman, J., Zhou, T., Chen, Z., Yuen, K.-Y., Kwong, P. D., Sodroski, J. G., Yin, M. T., Sheng, Z., Huang, Y., Shapiro, L., Ho, D. D. Nature 2020, 584, 450–456.

13. Washington, N. L., Gangavarapu, K., Zeller, M., Bolze, A., Cirulli, E. T., Schiabor Barrett, K. M., Larsen, B. B., Anderson, C., White, S., Cassens, T., Jacobs, S., Levan, G., Nguyen, J., Ramirez, J. M., Rivera-Garcia, C., Sandoval, E., Wang, X., Wong, D., Spencer, E., Robles-Sikisaka, R., Kurzban, E., Hughes, L. D., Deng, X., Wang, C., Servellita, V., Valentine, H., De Hoff, P., Seaver, P., Sathe, S., Gietzen, K., Sickler, B., Antico, J., Hoon, K., Liu, J., Harding, A., Bakhtar, O., Basler, T., Austin, B., MacCannell, D., Isaksson, M., Febbo, P. G., Becker, D., Laurent, M., McDonald, E., Yeo, G. W., Knight, R., Laurent, L. C., de Feo, E., Worobey, M., Chiu, C. Y., Suchard, M. A., Lu, J. T., Lee, W., Andersen, K. G. Cell 2021.

14. Humphrey, W., Dalke, A., Schulten, K. J. Mol. Graph. 1996, 14, 33-8, 27-8.

15. Pinto, D., Park, Y. J., Beltramello, M., Walls, A. C., Tortorici, M. A., Bianchi, S., Jaconi, S., Culap, K., Zatta, F., De Marco, A., Peter, A., Guarino, B., Spreafico, R., Cameroni, E., Case, J. B., Chen, R. E., Havenar-Daughton, C., Snell, G., Telenti, A., Virgin, H. W., Lanzavecchia, A., Diamond, M. S., Fink, K., Veesler, D., Corti, D. bioRxiv 2020.

16. Woo, H., Park, S.-J., Choi, Y. K., Park, T., Tanveer, M., Cao, Y., Kern, N. R., Lee, J., Yeom, M. S., Croll, T. I., Seok, C., Im, W. J. Phys. Chem. B 2020, 124, 7128–7137.

17. Choi, Y. K., Cao, Y., Frank, M., Woo, H., Park, S.-J., Yeom, M. S., Croll, T. I., Seok, C., Im, W. J. Chem. Theory Comput. 2021, 17, 2479–2487.

18. Ko, J., Park, H., Seok, C. BMC Bioinformatics 2012, 13, 198.

19. Ko, J., Lee, D., Park, H., Coutsias, E. A., Lee, J., Seok, C. Nucleic Acids Res. 2011, 39, W210–W214.

20. Park, T., Baek, M., Lee, H., Seok, C. J. Comput. Chem. 2019, 40, 2413–2417.

21. Croll, T. I. Acta Crystallogr. D 2018, 74, 519–530.

22. Jo, S., Song, K. C., Desaire, H., MacKerell, A. D., Jr., Im, W. J. Comput. Chem. 2011, 32, 3135–41.

23. Park, S. J., Lee, J., Patel, D. S., Ma, H., Lee, H. S., Jo, S., Im, W. Bioinformatics 2017, 33, 3051–3057.

24. Park, S. J., Lee, J., Qi, Y., Kern, N. R., Lee, H. S., Jo, S., Joung, I., Joo, K., Lee, J., Im, W. Glycobiology 2019, 29, 320–331.

25. Yuan, A., Jo, S., Kim, T., Im, W. PLoS ONE 2007, 2, e880.

26. Jo, S., Lim, J. B., Klauda, J. B., Im, W. Biophys. J. 2009, 97, 50–58.

27. Wu, E. L., Cheng, X., Jo, S., Rui, H., Song, K. C., Dávila-Contreras, E. M., Qi, Y., Lee, J., Monje-Galvan, V., Venable, R. M., Klauda, J. B., Im, W. J. Comput. Chem. 2014, 35, 1997–2004.

28. Lee, J., Patel, D. S., Ståhle, J., Park, S.-J., Kern, N. R., Kim, S., Lee, J., Cheng, X., Valvano, M. A., Holst, O., Knirel, Y. A., Qi, Y., Jo, S., Klauda, J. B., Widmalm, G., Im, W. J. Chem. Theory Comput. 2018, 15, 775–786.

29. Huang, J., Rauscher, S., Nawrocki, G., Ran, T., Feig, M., de Groot, B. L., Grubmüller, H., MacKerell, A. D. Nat. Methods 2016, 14, 71–73.

30. Guvench, O., Greene, S. N., Kamath, G., Brady, J. W., Venable, R. M., Pastor, R. W., Mackerell, A. D. J. Comput. Chem. 2008, 29, 2543–2564.

31. Guvench, O., Hatcher, E., Venable, R. M., Pastor, R. W., MacKerell, A. D. J. Chem. Theory Comput. 2009, 5, 2353–2370.

32. Hatcher, E., Guvench, O., MacKerell, A. D. J. Phys. Chem. B 2009, 113, 12466–12476.

33. Jorgensen, W. L., Chandrasekhar, J., Madura, J. D., Impey, R. W., Klein, M. L. J. Chem. Phys. 1983, 79, 926–935.

34. Steinbach, P. J., Brooks, B. R. J. Comput. Chem. 1994, 15, 667–683.

35. Essmann, U., Perera, L., Berkowitz, M. L., Darden, T., Lee, H., Pedersen, L. G. J. Chem. Phys. 1995, 103, 8577–8593.

36. Nawrocki, G., Im, W., Sugita, Y., Feig, M. Proc. Natl. Acad. Sci. U.S.A. 2019, 116, 24562–24567.

37. Nawrocki, G., Wang, P.-h., Yu, I., Sugita, Y., Feig, M. J. Phys. Chem. B 2017, 121, 11072–11084.

38. Jo, S., Kim, T., Iyer, V. G., Im, W. J. Comput. Chem. 2008, 29, 1859–65.

39. Lee, J., Cheng, X., Swails, J. M., Yeom, M. S., Eastman, P. K., Lemkul, J. A., Wei, S., Buckner, J., Jeong, J. C., Qi, Y., Jo, S., Pande, V. S., Case, D. A., Brooks, C. L., MacKerell, A. D., Klauda, J. B., Im, W. J. Chem. Theory Comput. 2015, 12, 405–413.

40. Lee, J., Hitzenberger, M., Rieger, M., Kern, N. R., Zacharias, M., Im, W. J. Chem. Phys. 2020, 153.

41. Van Der Spoel, D., Lindahl, E., Hess, B., Groenhof, G., Mark, A. E., Berendsen, H. J. C. J. Comput. Chem. 2005, 26, 1701–1718.

42. Hess, B., Bekker, H., Berendsen, H. J. C., Fraaije, J. G. E. M. J. Comput. Chem. 1997, 18, 1463–1472.

43. Nosé, S. Mol. Phys. 1984, 52, 255–268.

44. Hoover, W. G. Phys. Rev. A 1985, 31, 1695–1697.

45. Parrinello, M., Rahman, A. J. Appl. Phys. 1981, 52, 7182–7190.

46. Nosé, S., Klein, M. L. Mol. Phys. 1983, 50, 1055–1076.

47. Hopkins, C. W., Le Grand, S., Walker, R. C., Roitberg, A. E. J. Chem. Theory Comput. 2015, 11, 1864–1874.

48. Li, Q., Wu, J., Nie, J., Zhang, L., Hao, H., Liu, S., Zhao, C., Zhang, Q., Liu, H., Nie, L., Qin, H., Wang, M., Lu, Q., Li, X., Sun, Q., Liu, J., Zhang, L., Li, X., Huang, W., Wang, Y. Cell 2020, 182, 1284–1294.e9.

49. Cao, W., Dong, C., Kim, S., Hou, D., Tai, W., Du, L., Im, W., Zhang, X. F. Biophys. J. 2021, 120, 1011–1019.

50. Casalino, L., Gaieb, Z., Goldsmith, J. A., Hjorth, C. K., Dommer, A. C., Harbison, A. M., Fogarty, C. A., Barros, E. P., Taylor, B. C., McLellan, J. S., Fadda, E., Amaro, R. E. ACS Cent. Sci. 2020, 6, 1722–1734.

