## Supporting Information for "Dynamic Interactions of Fully Glycosylated SARS-CoV-2 Spike Protein with Various Antibodies"

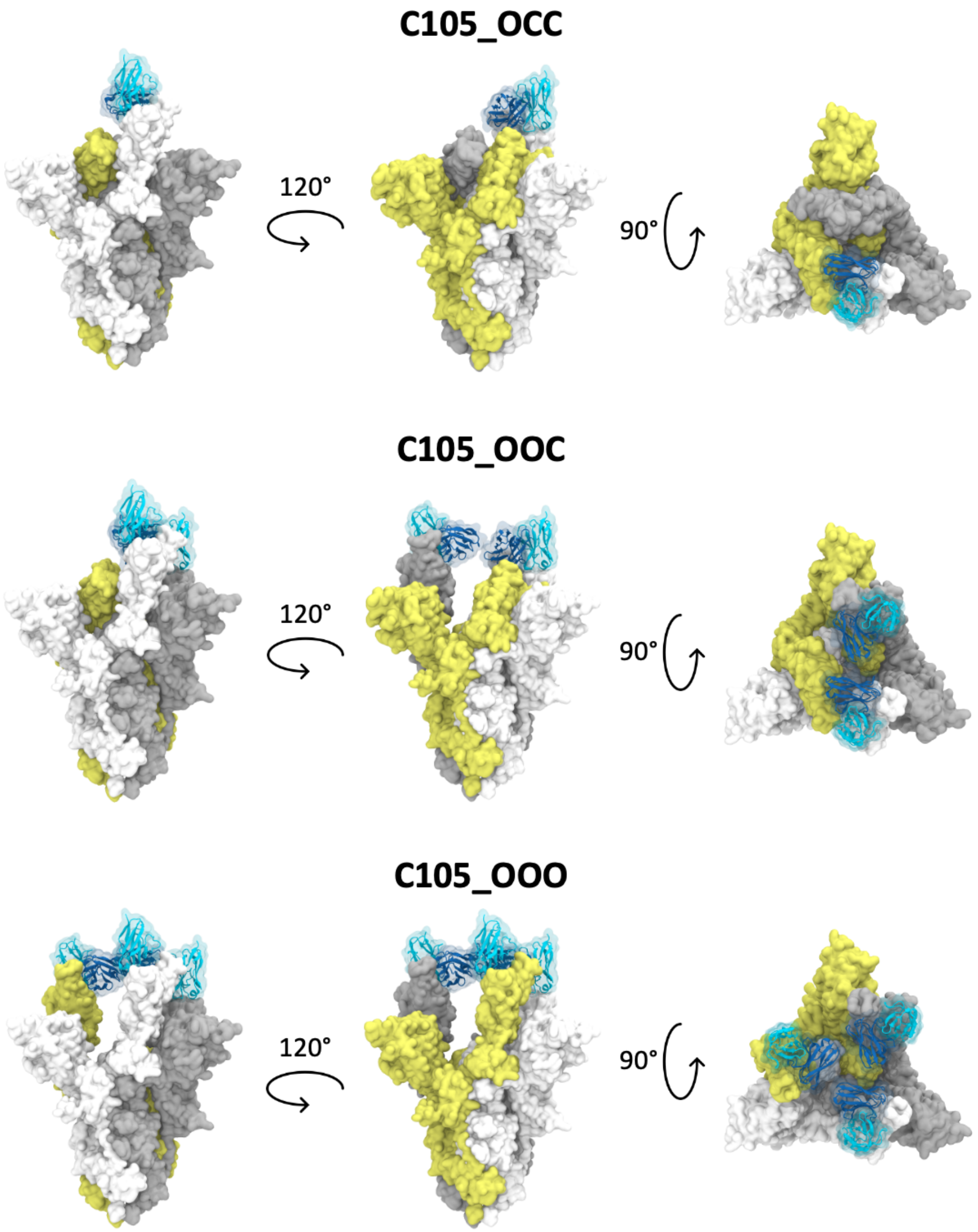


**Figure S1. Illustration of C105_OCC, C105_OOC and C105_OOO.** The three protomers of S protein are shown in white, gray, and yellow respectively, and the antibodies are shown in blue.

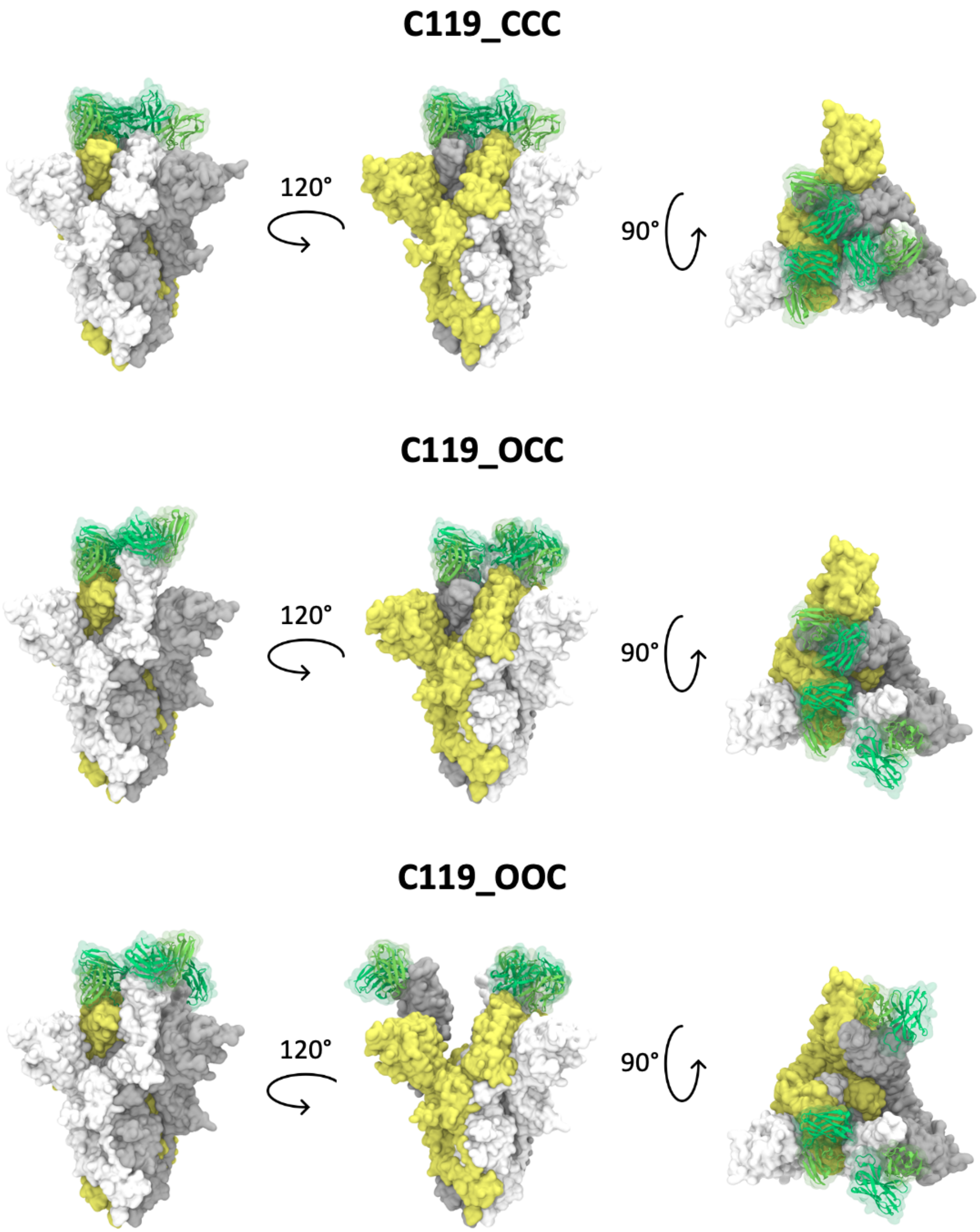


**Figure S2. Illustration of C119_CCC, C119_OCC and C119_OOC.** The three protomers of S protein are shown in white, gray, and yellow respectively, and the antibodies are shown in green.

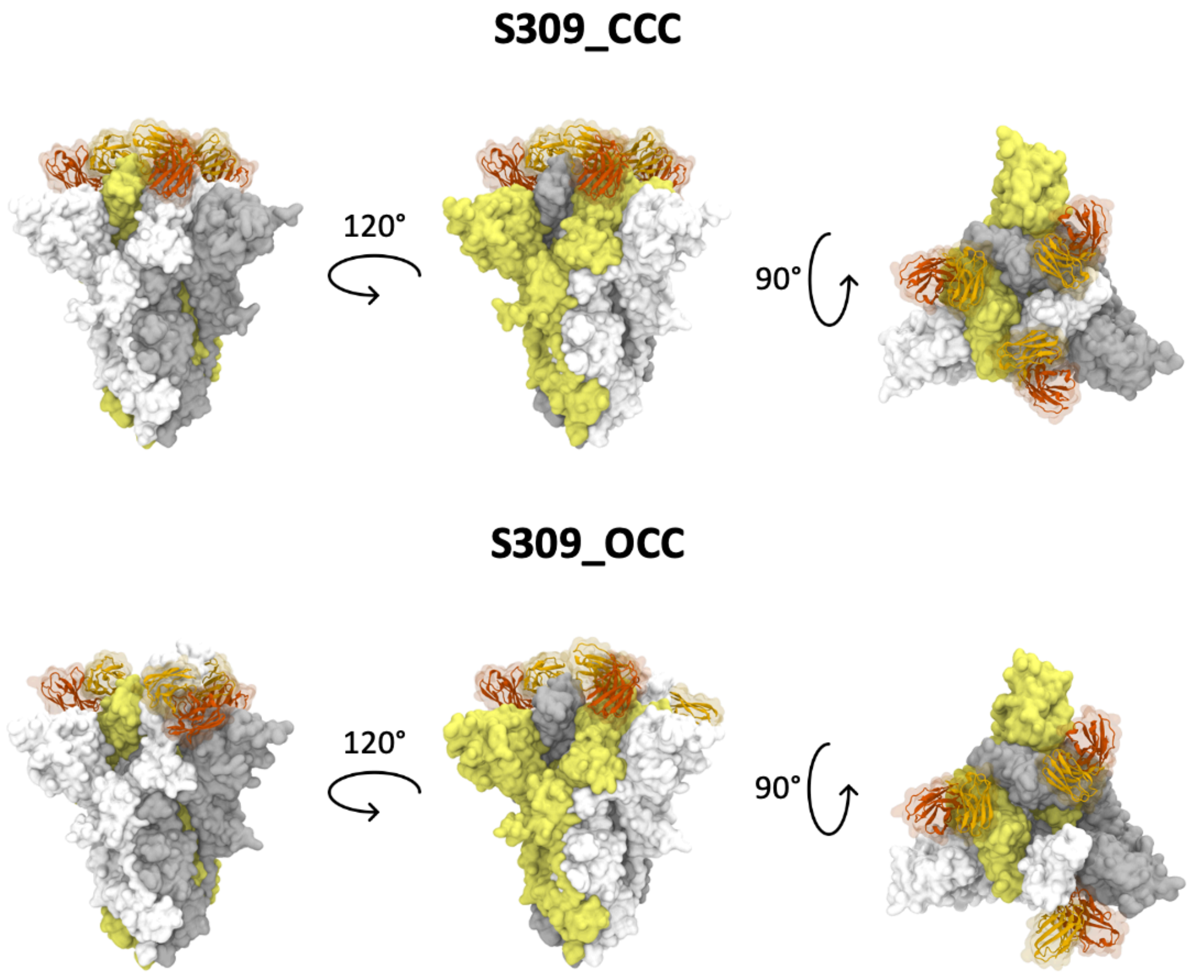


**Figure S3. Illustration of S309_CCC, and S309_OCC.** The three protomers of S protein are shown in white, gray, and yellow respectively, and the antibodies are shown in orange.

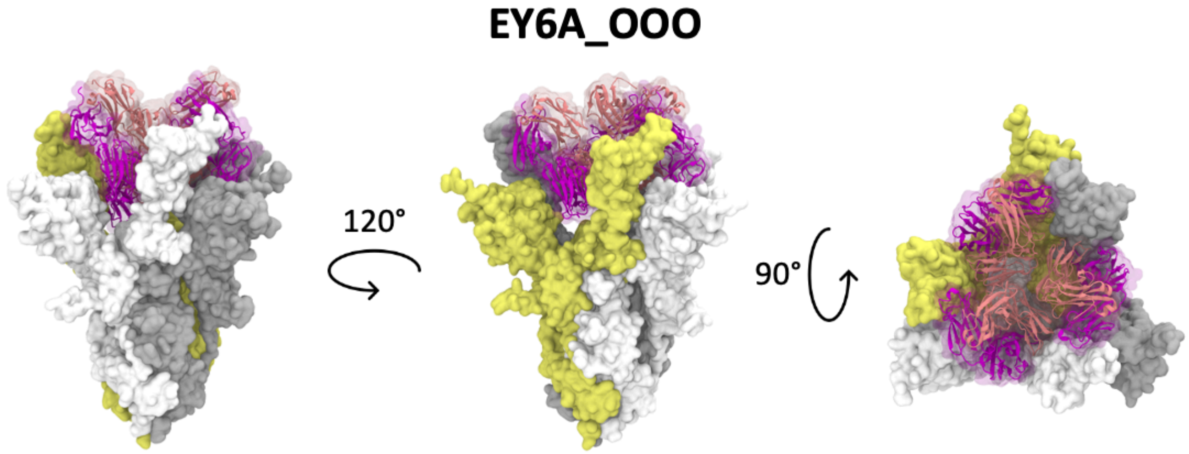


**Figure S4. Illustration of EY6A_OOO.** The three protomers of S protein are shown in white, gray, and yellow respectively, and the antibodies are shown in magenta.

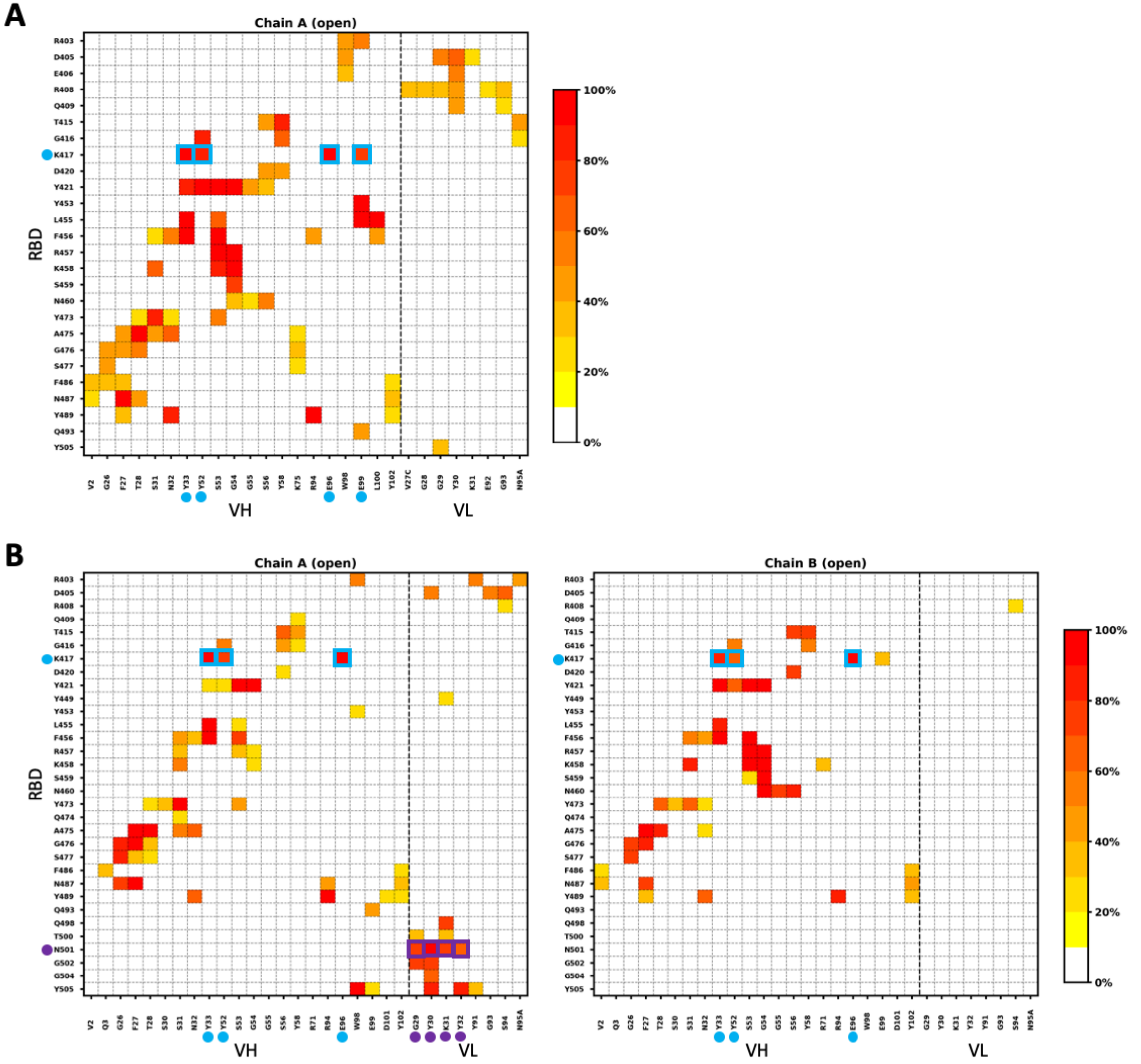


**Figure S5. The frequency of interacting residue pairs in C105 systems.** (A) C105_OCC, (B) C105_OOC, and (C) C105_OOO.

C

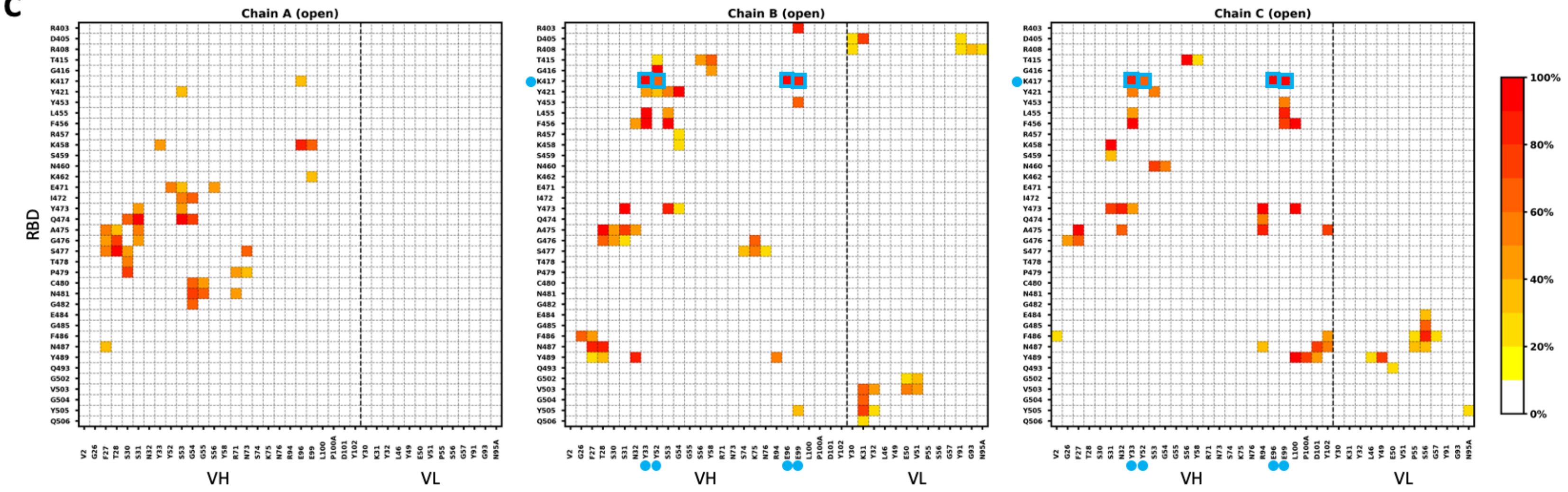


**Figure S5 (Continued). The frequency of interacting residue pairs in C105 systems.** (A) C105_OCC, (B) C105_OOC, and (C) C105_OOO.

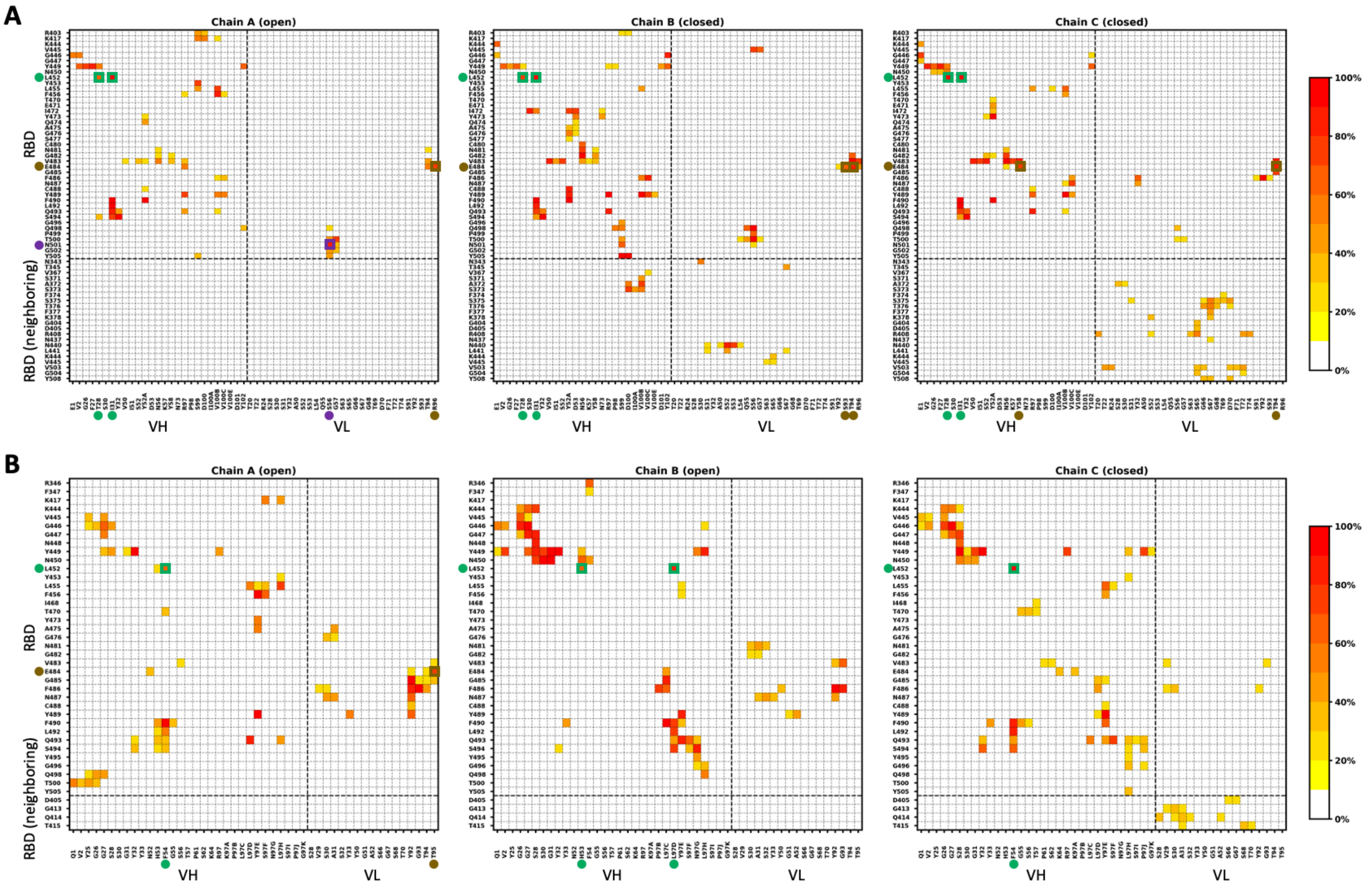


**Figure S6. The frequency of interacting residue pairs in C002 systems.** (A) C002_OCC and (B) C002_OOC.

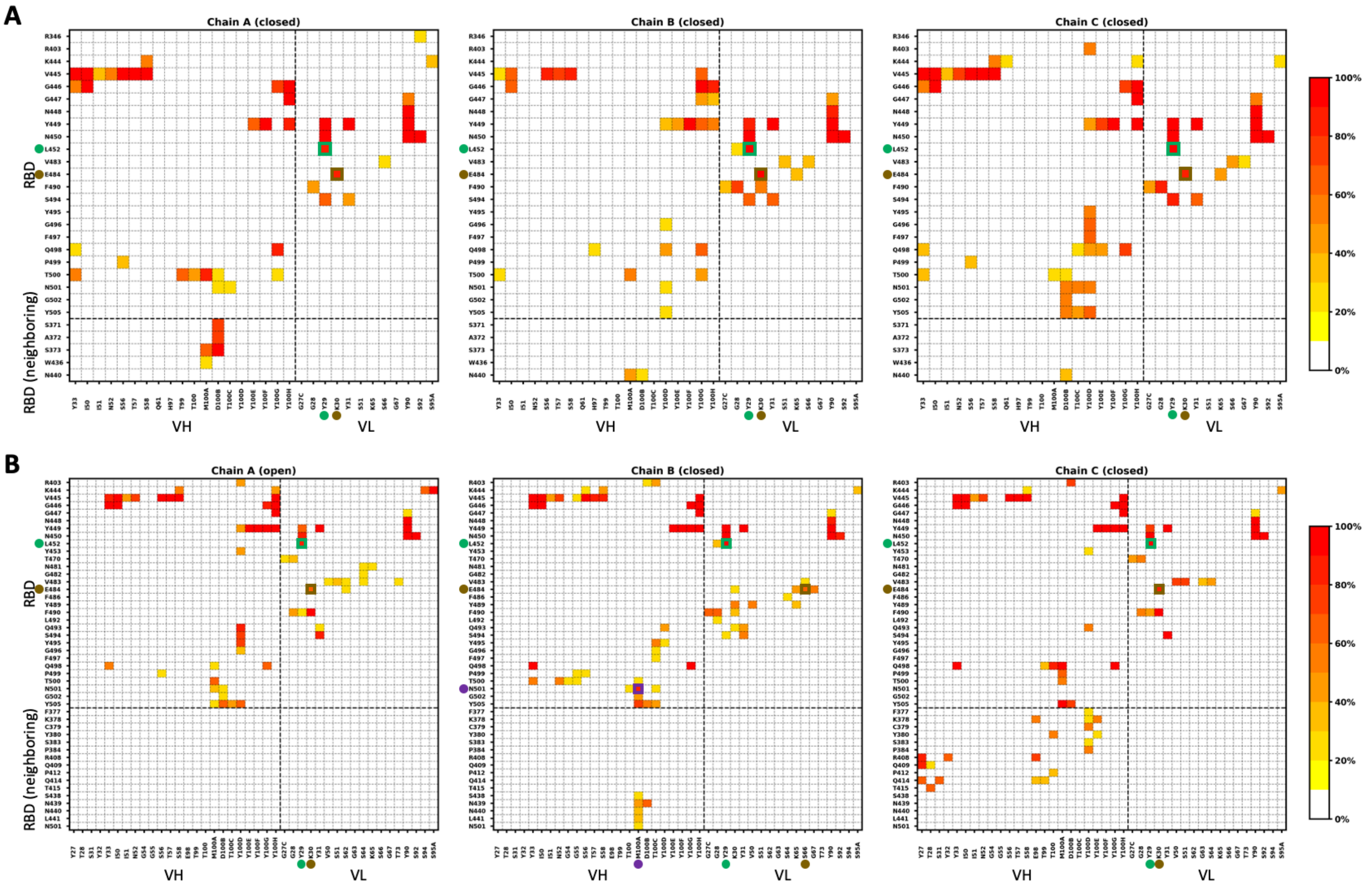


**Figure S7. The frequency of interacting residue pairs in C119 systems.** (A) C119_CCC, (B) C119_OCC, and (C) C119_OOC.

**C**

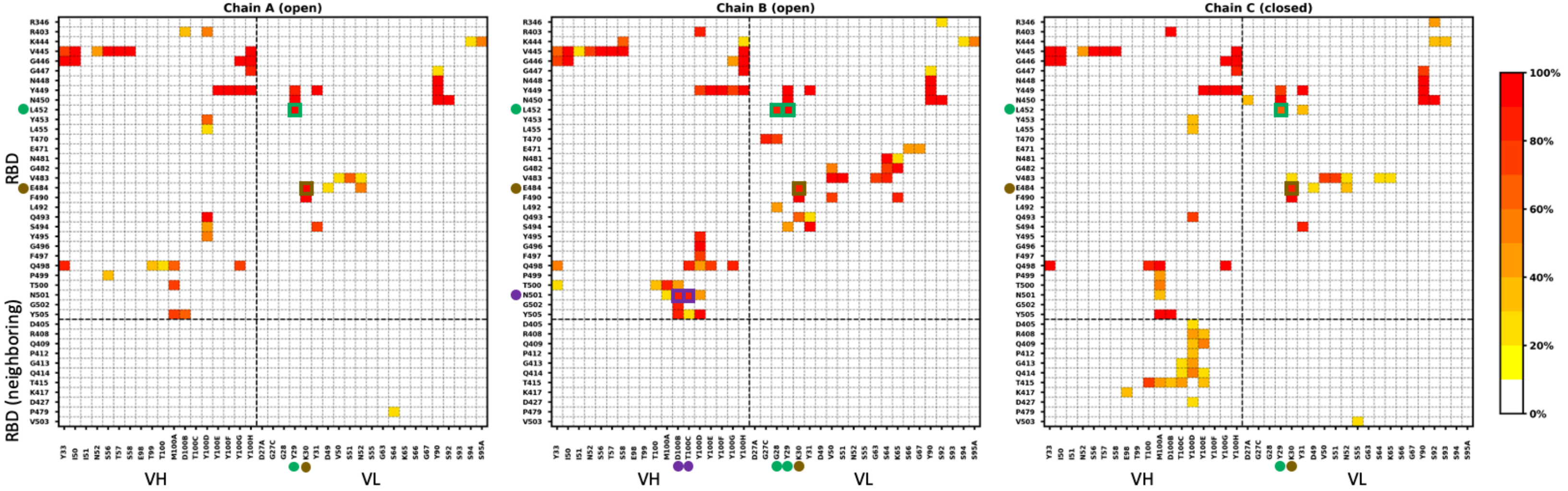


**Figure S7 (Continued). The frequency of interacting residue pairs in C119 systems.** (A) C119_CCC, (B) C119_OCC, and (C) C119_OOC.

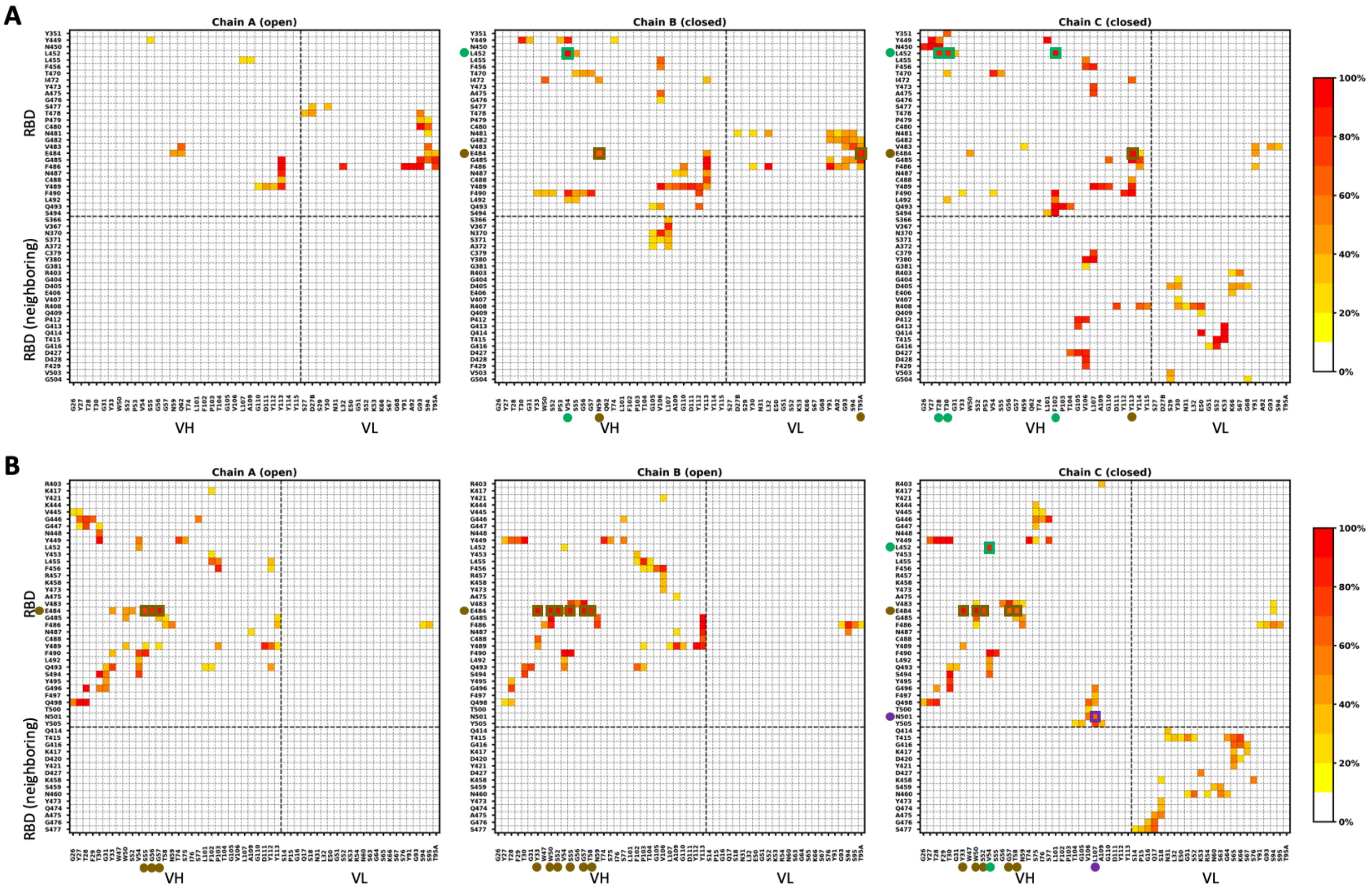


**Figure S8. The frequency of interacting residue pairs in C121 systems.** (A) C121_OCC and (B) C121_OOC.

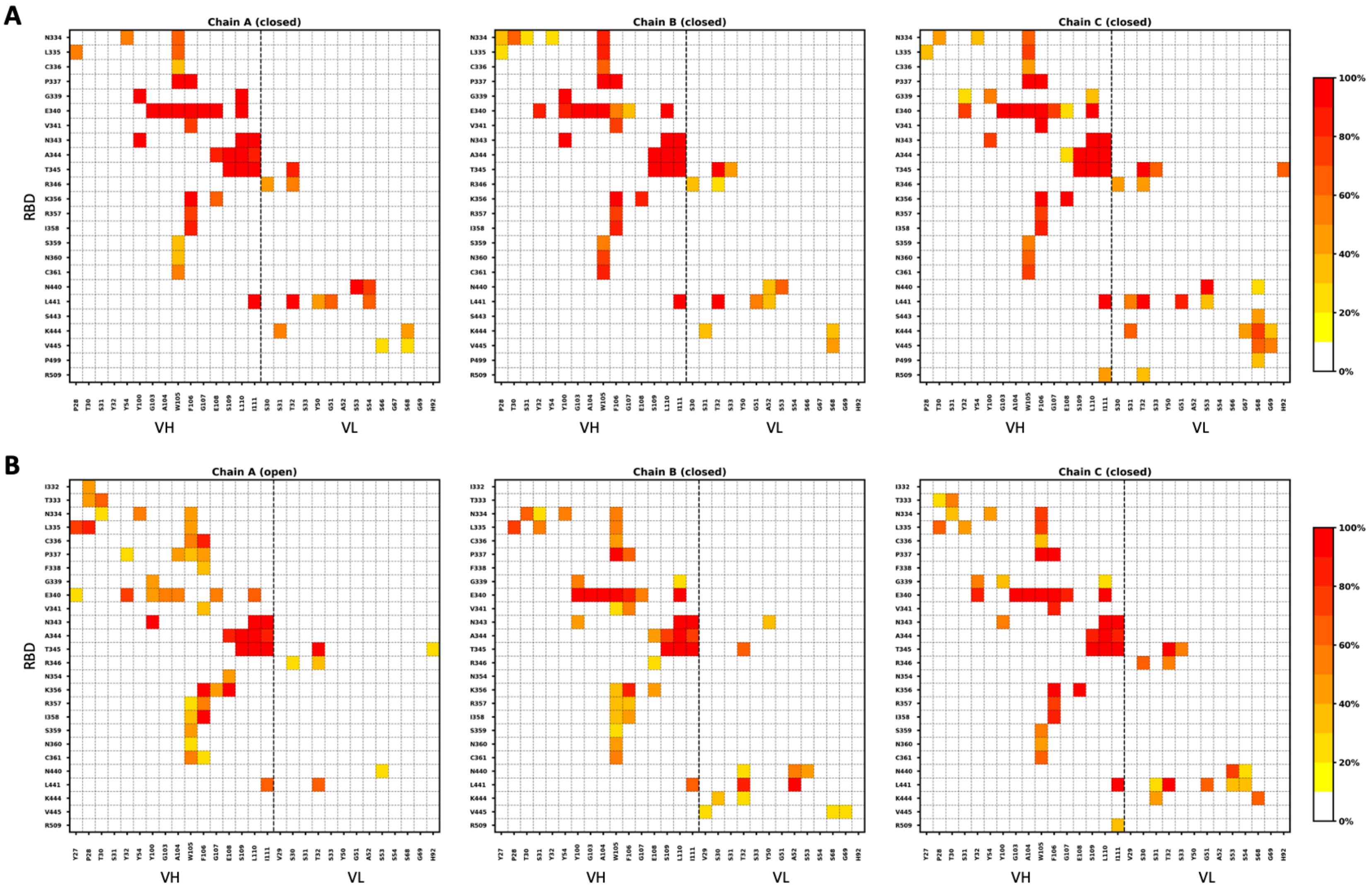


**Figure S9. The frequency of interacting residue pairs in S309 systems.** (A) S309_CCC and (B) S309_OCC.

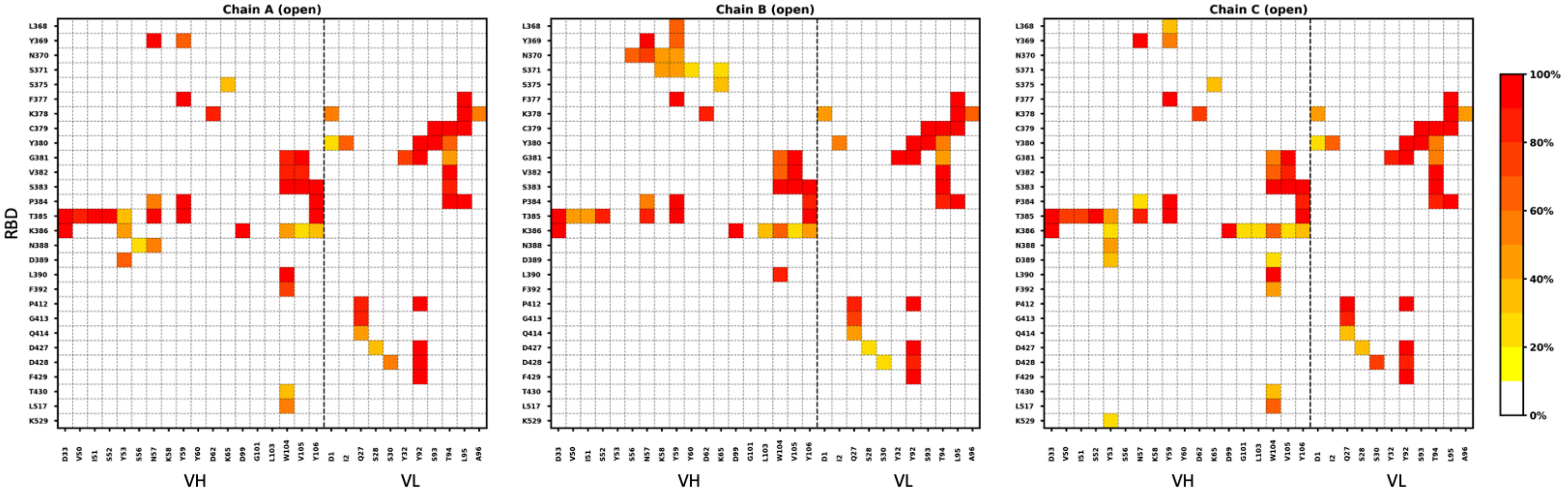


**Figure S10. The frequency of interacting residue pairs in EY6A_OOO.**

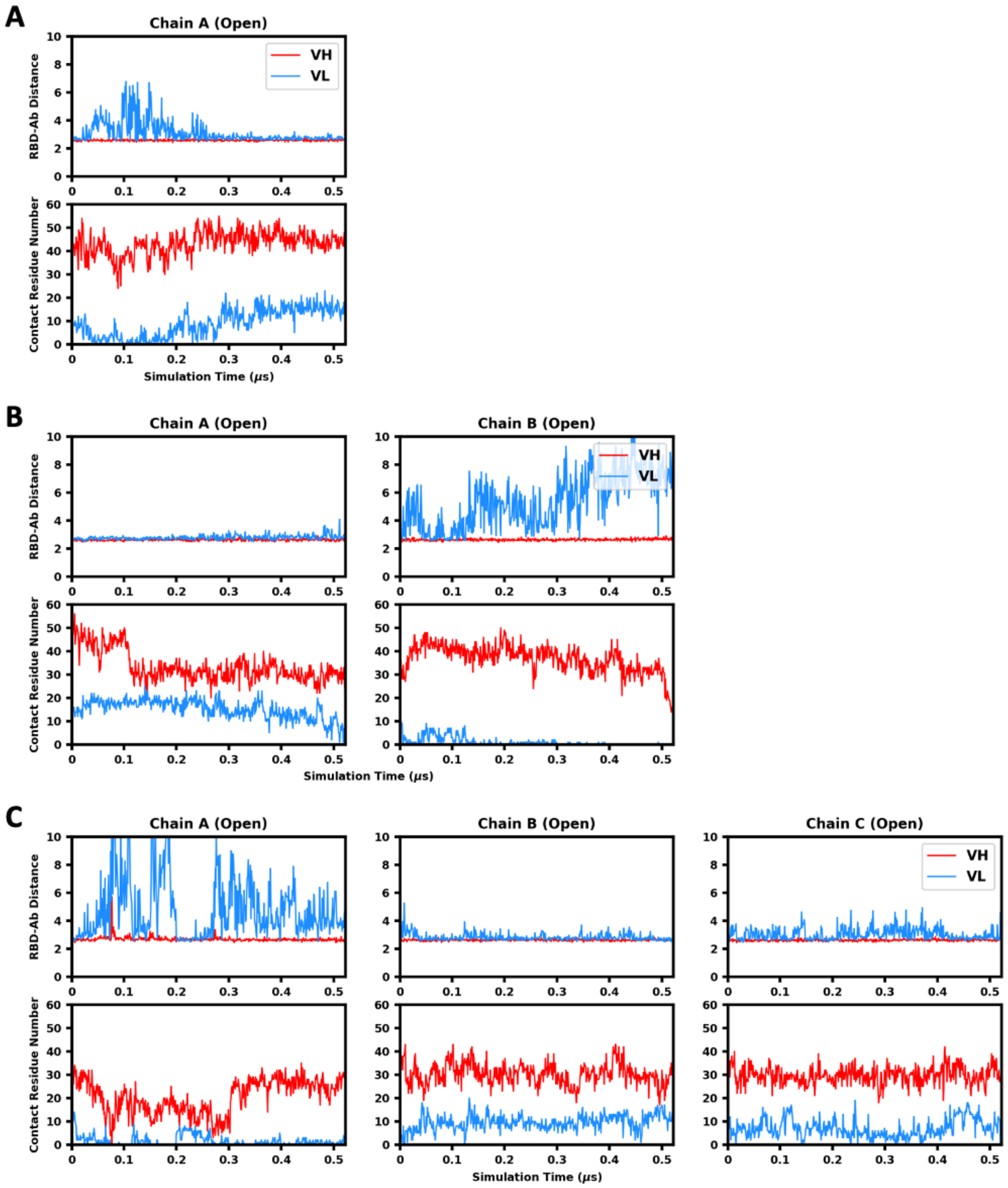


**Figure S11. RBD-antibody distance and contact residue number in C105 systems.** (A) C105_OCC, (B) C105_OOC, and (C) C105_OOO.

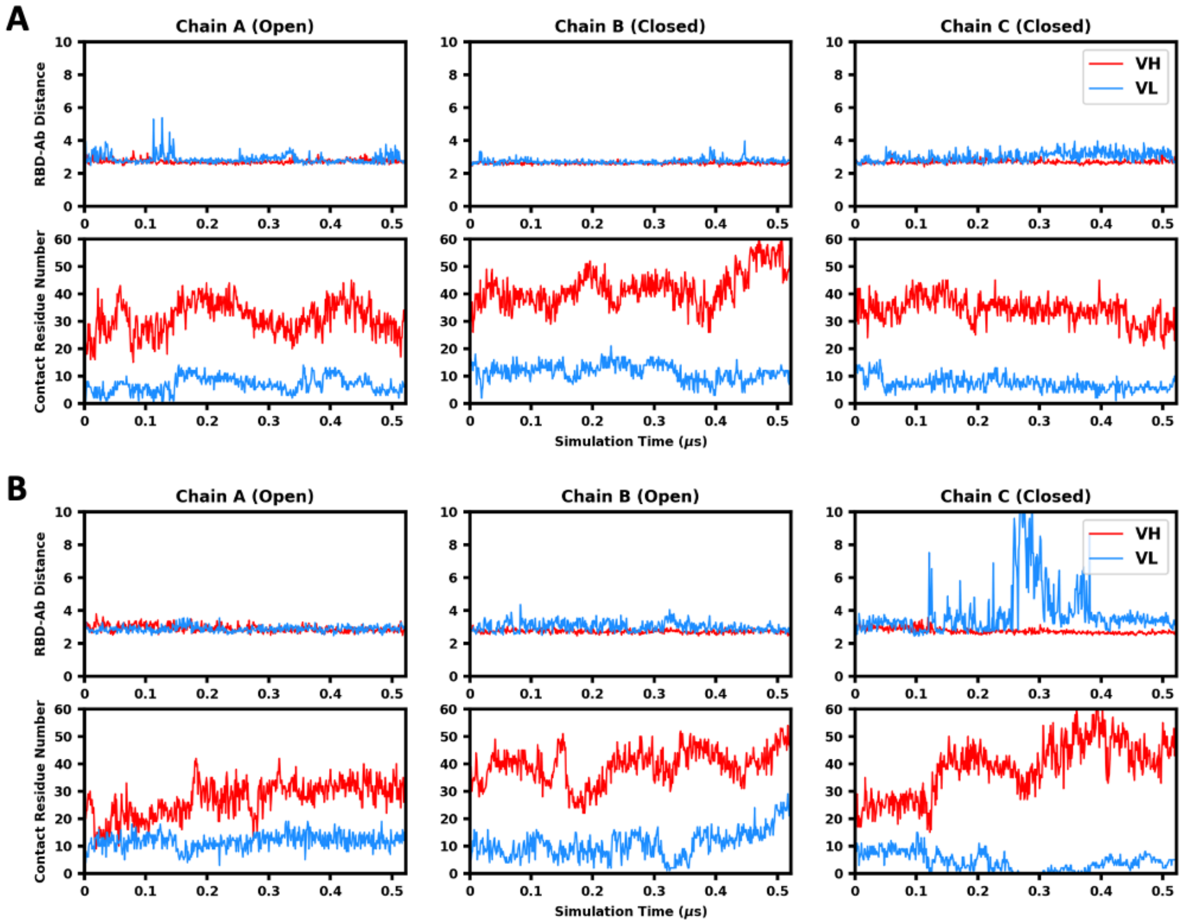


**Figure S12. RBD-antibody distance and contact residue number in C002 systems.** (A) C002_OCC and (B) C002_OOC.

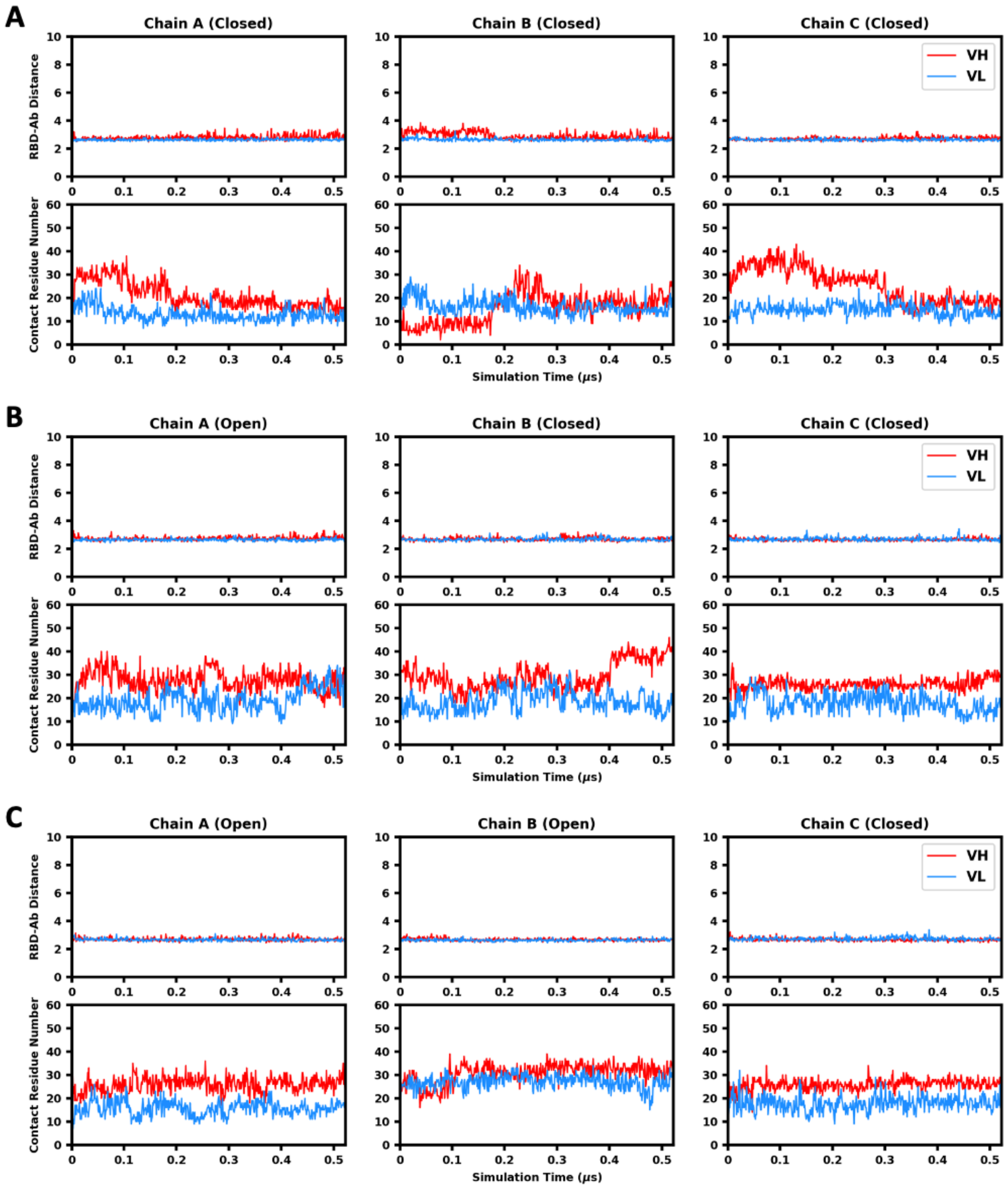


**Figure S13. RBD-antibody distance and contact residue number in C119 systems.** (A) C119_CCC, (B) C119_OCC, and (C) C119_OOC.

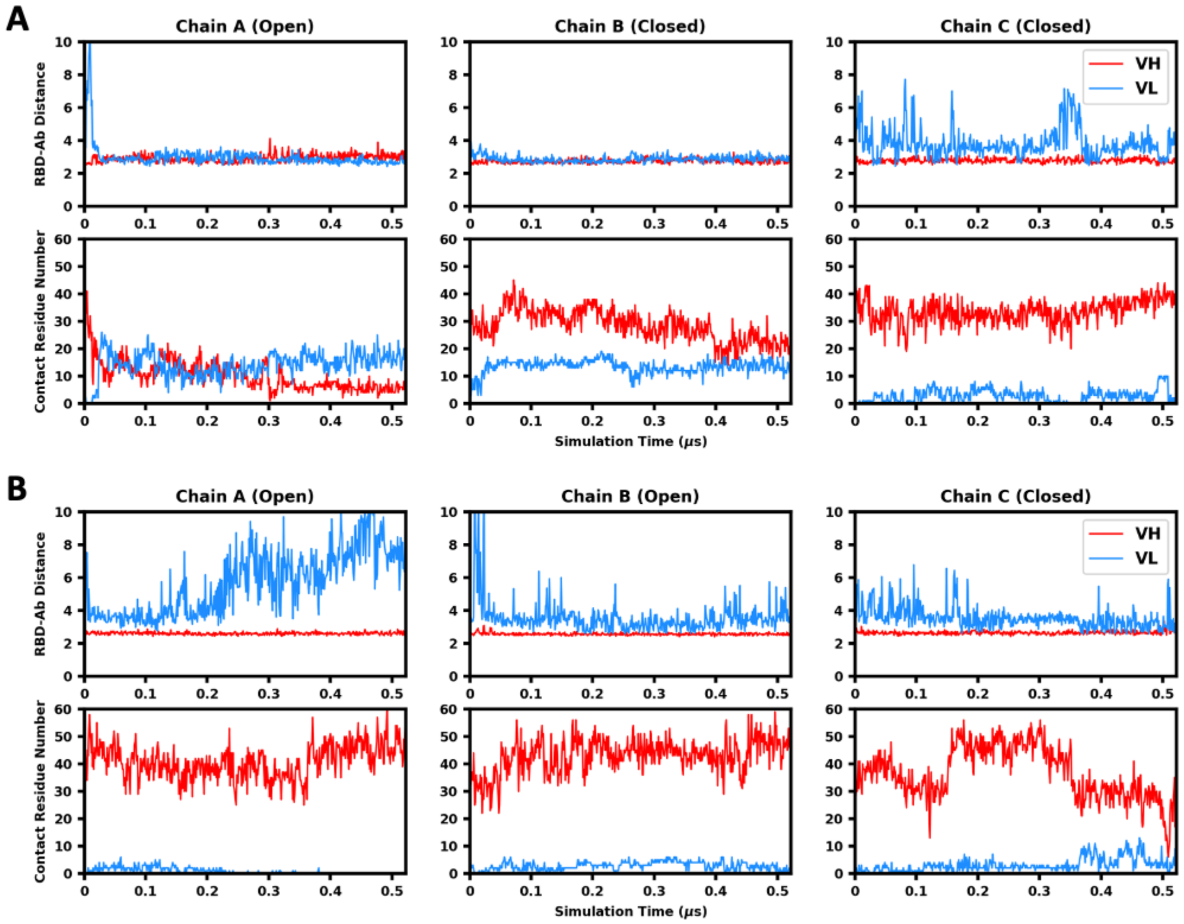


**Figure S14. RBD-antibody distance and contact residue number in C121 systems.** (A) C121_OCC and (B) C121_OOC.

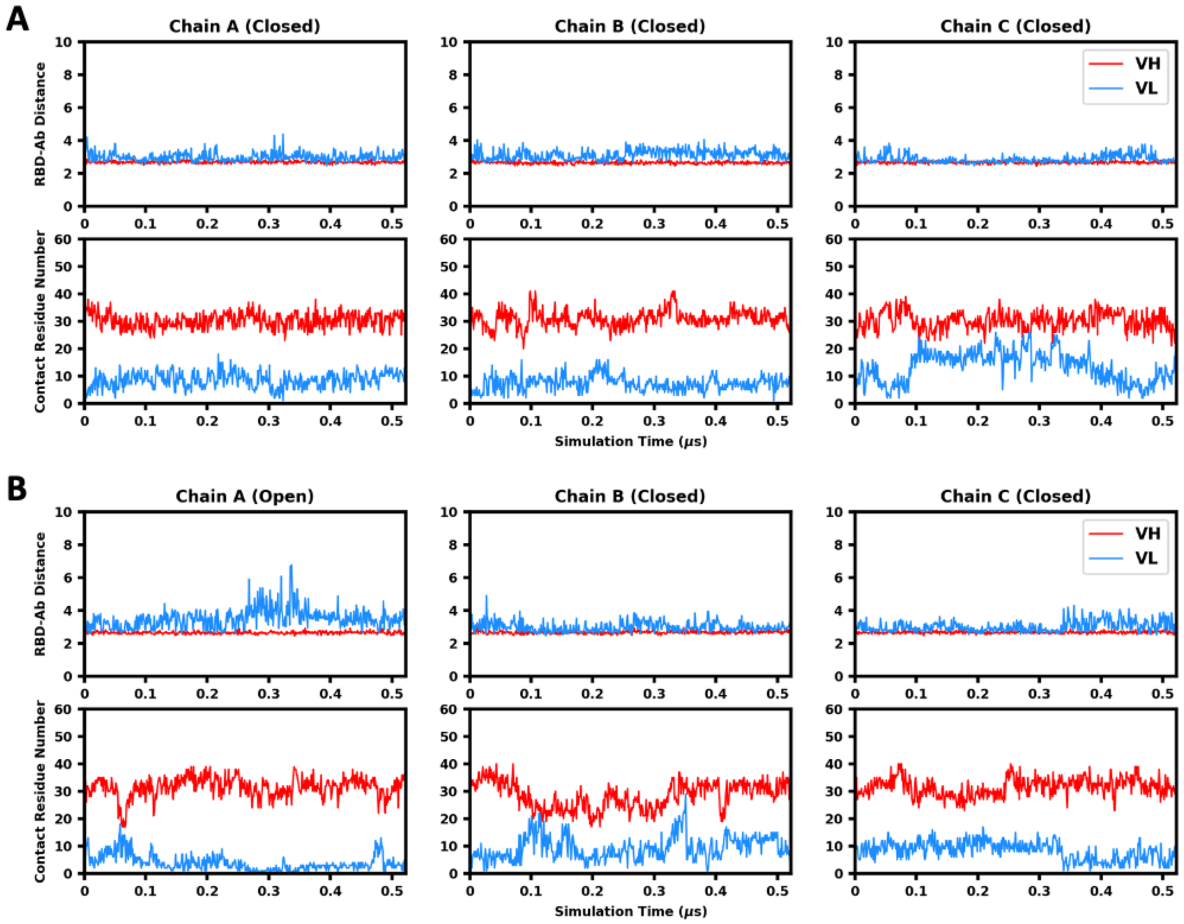


**Figure S15. RBD-antibody distance and contact residue number in S309 systems.** (A) S309_CCC and (B) S309_OCC.

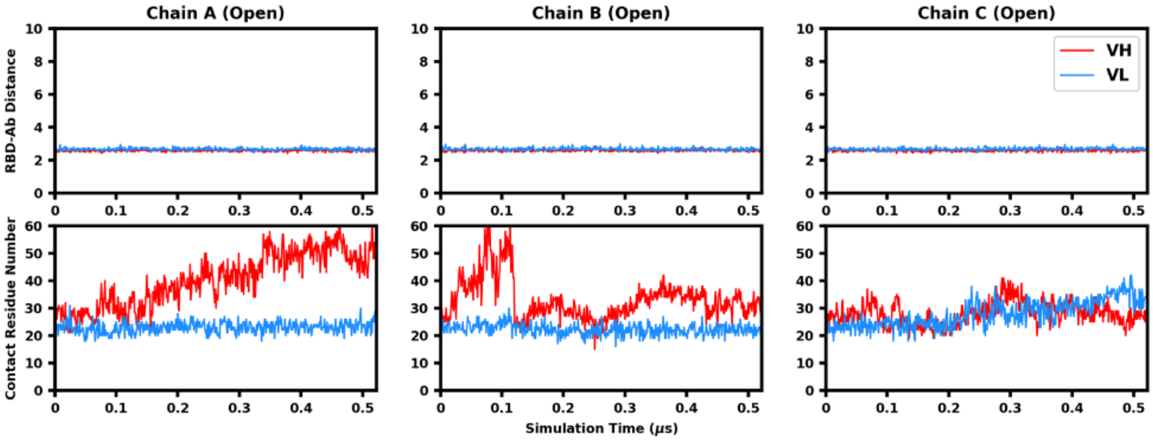


**Figure S16. RBD-antibody distance and contact residue number in EY6A_OOO.**

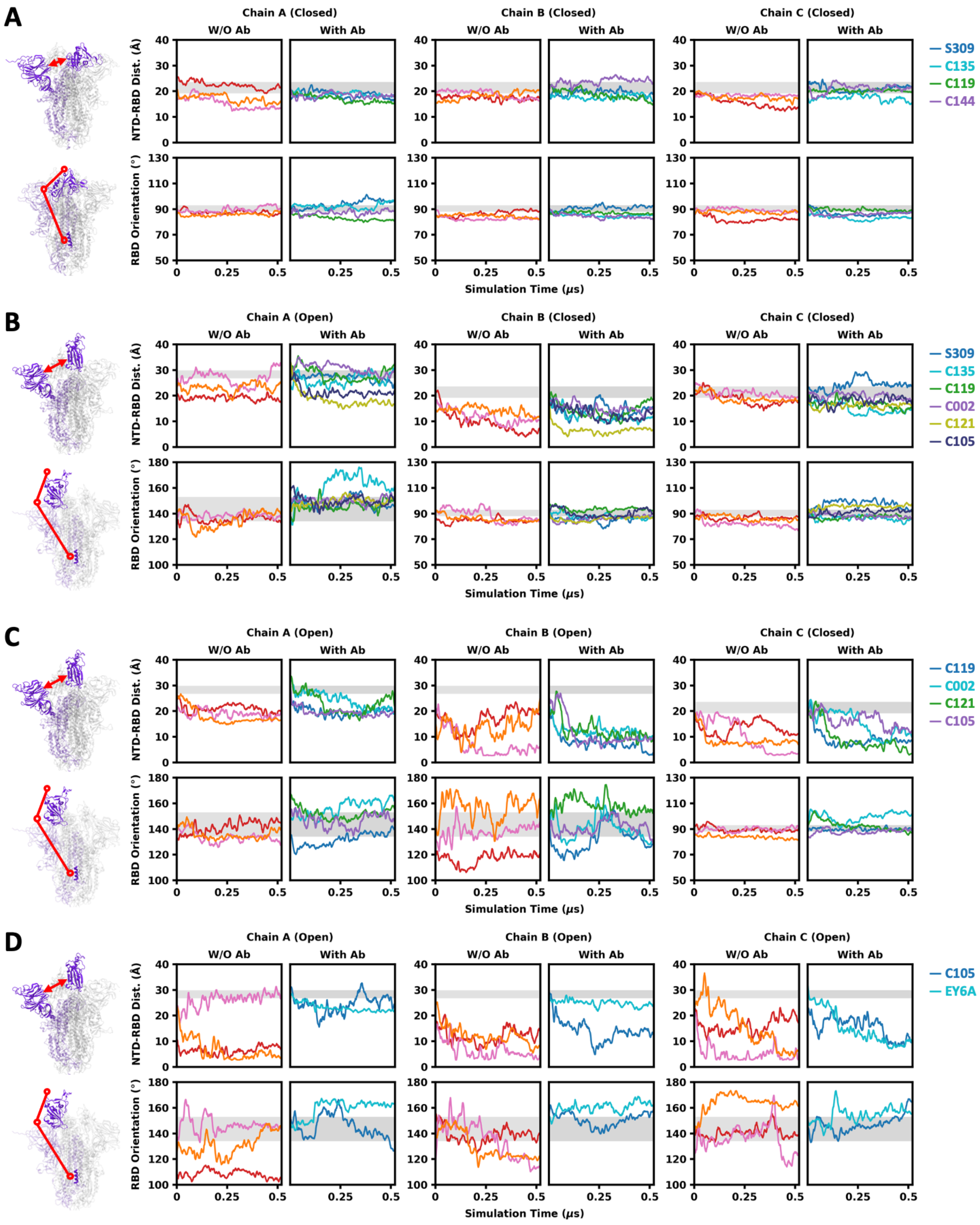


**Figure S17. NTD-RBD distance and RBD orientation in S-only and S-antibody complex systems.** (A) All closed, (B) one RBD open, (C) two RBDs open, and (D) all open. Three trajectories of S-only systems are shown in red, pink, and orange. The trajectories of S protein in complex with various antibodies are shown in the colors labeled on the right.

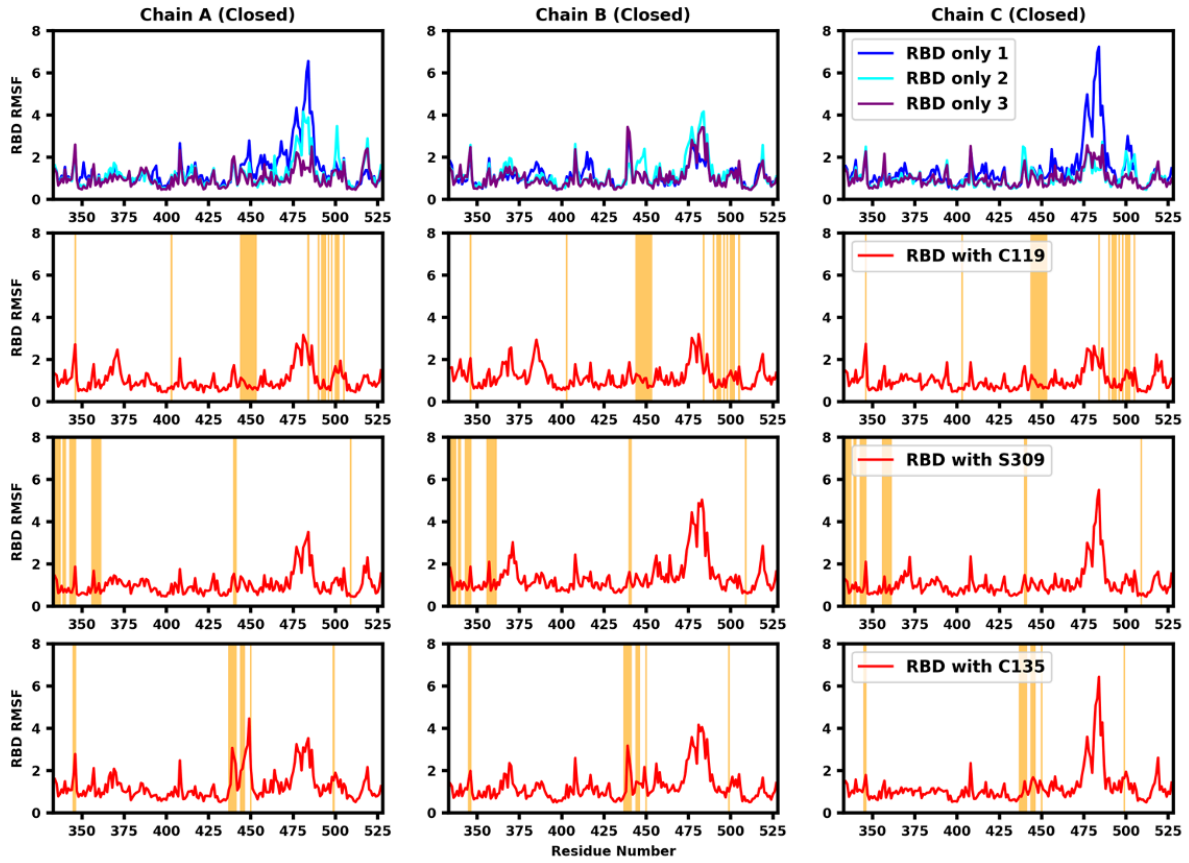


**Figure S18. RMSF of RBD in S-only and S-antibody complex systems with all three RBDs closed.** Three trajectories of S-only systems are shown in blue, cyan, and purple, and the trajectories of S-antibody systems are shown in red. The residues in the epitope of each antibody are shaded by orange regions.

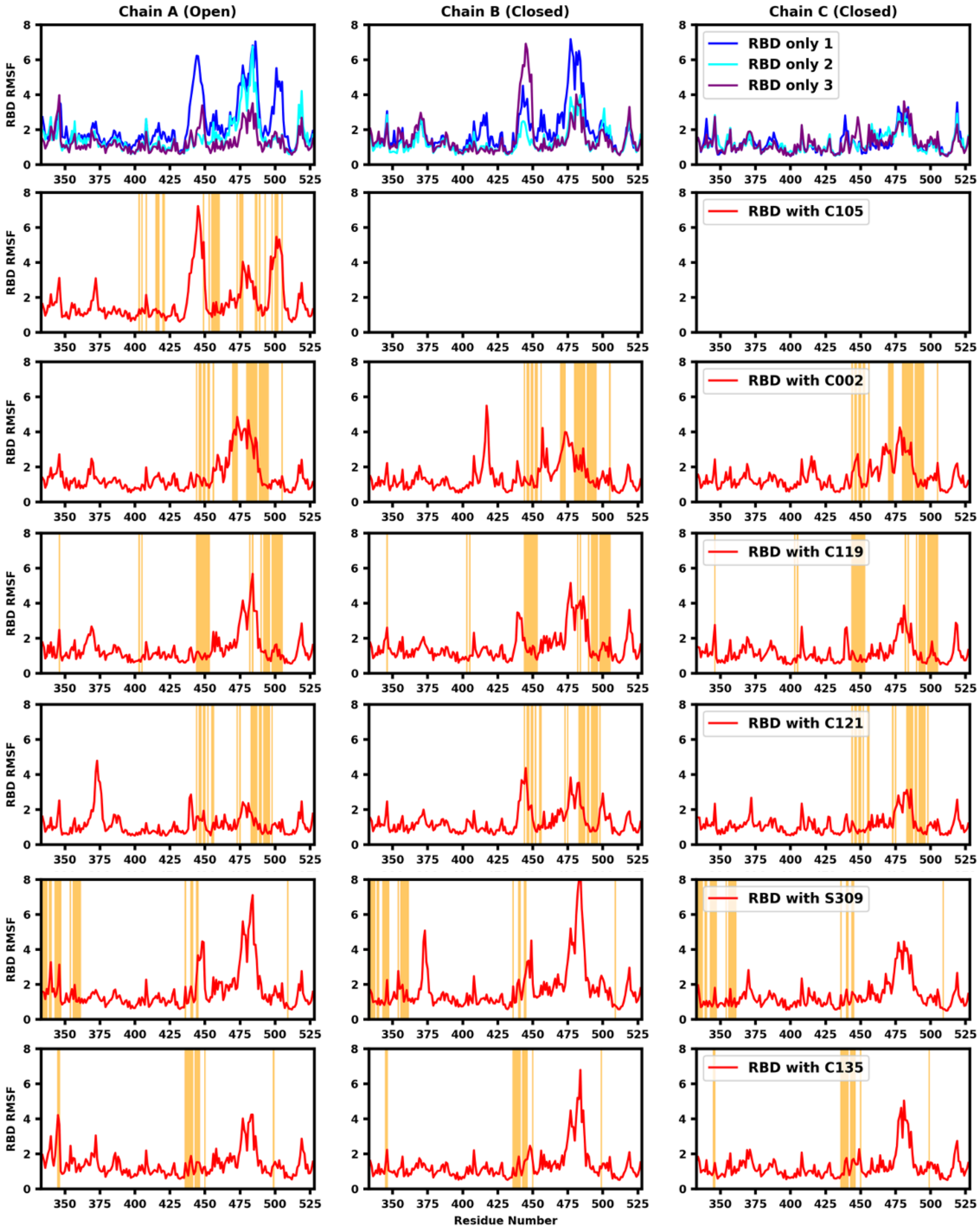


**Figure S19. RMSF of RBD in S-only and S-antibody complex systems with one RBD open and two RBDs closed.** Three trajectories of S-only systems are shown in blue, cyan, and purple, and the trajectories of S-antibody systems are shown in red. The residues in the epitope of each antibody are shaded by orange regions.

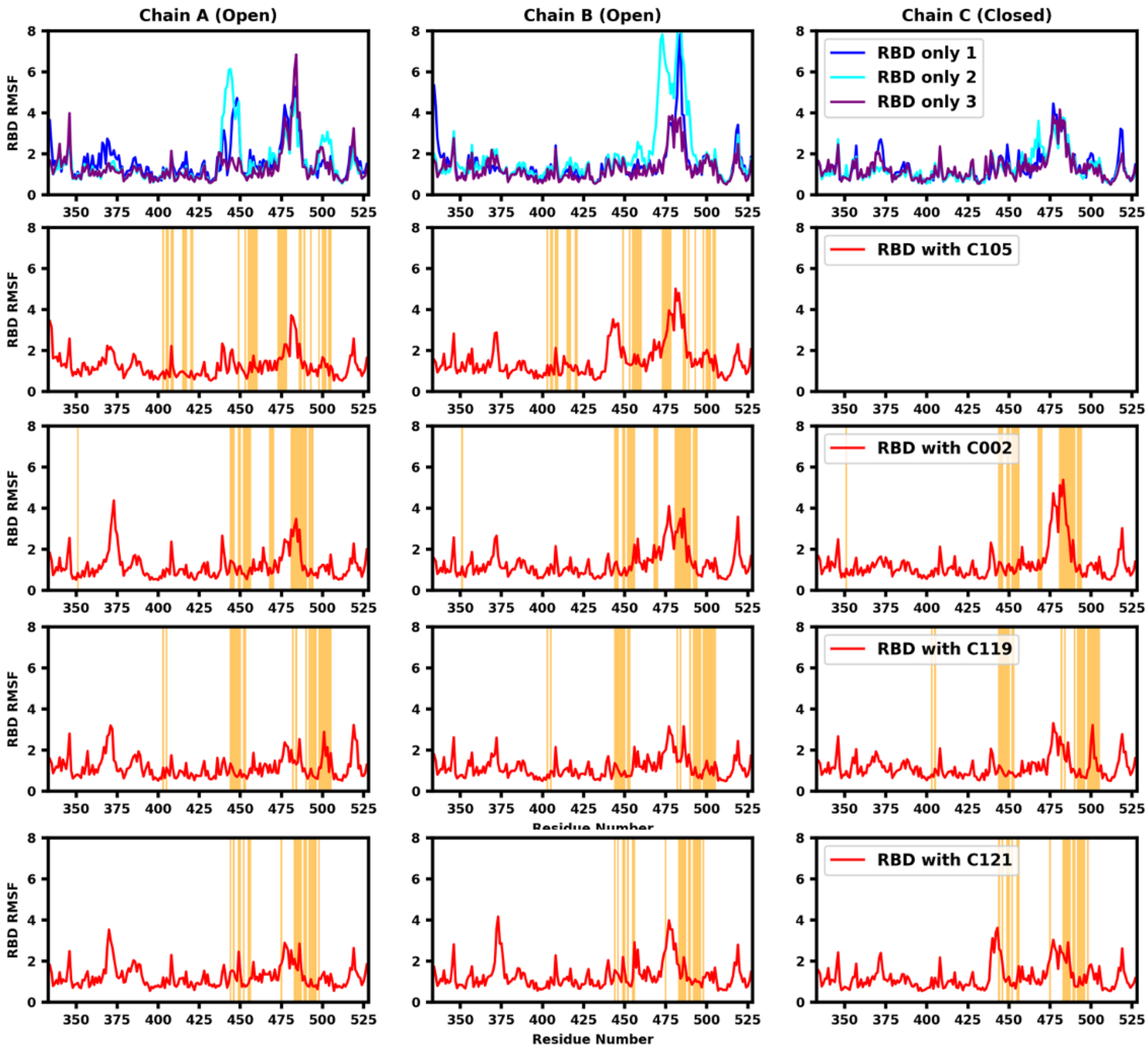


**Figure S20. RMSF of RBD in S-only and S-antibody complex systems with two RBDs open and one RBD closed.** Three trajectories of S-only systems are shown in blue, cyan, and purple, and the trajectories of S-antibody systems are shown in red. The residues in the epitope of each antibody are shaded by orange regions.

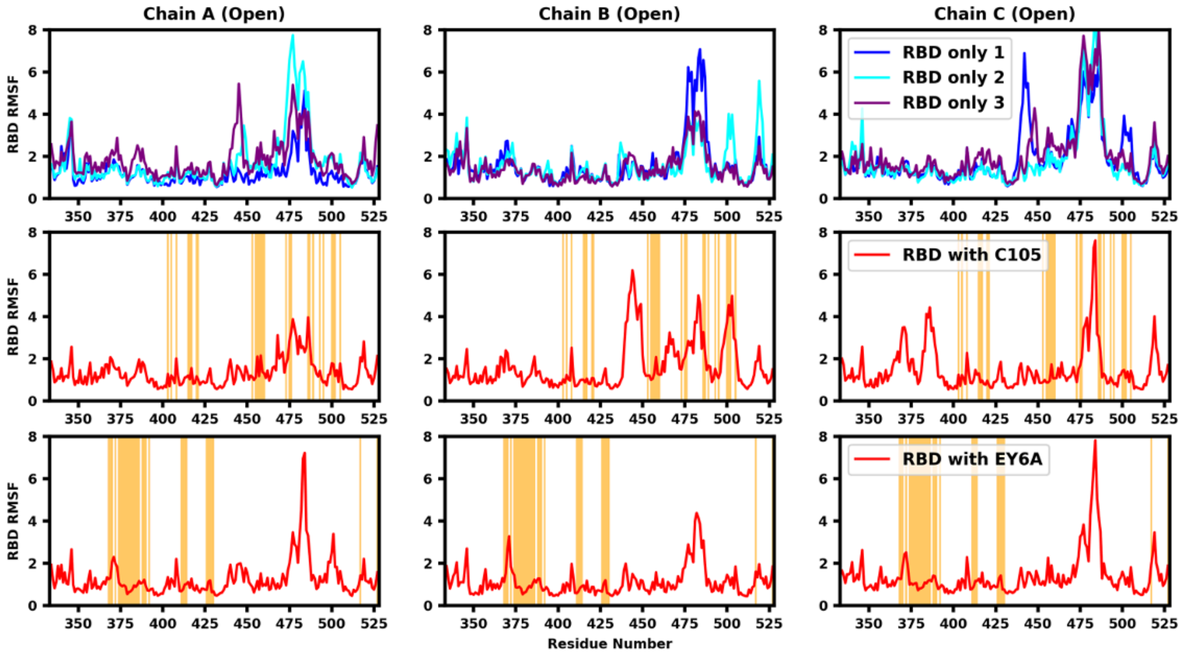


**Figure S21. RMSF of RBD in S-only and S-antibody complex systems with all three RBDs open.** Three trajectories of S-only systems are shown in blue, cyan, and purple, and the trajectories of S-antibody systems are shown in red. The residues in the epitope of each antibody are shaded by orange regions.
